# Detection of titanium nanoparticles in human, animal and infant formula milk

**DOI:** 10.1101/2024.10.03.616116

**Authors:** Camille Rivard, Nouzha Djebrani-Oussedik, Romane Cloix, Cathy Hue-Beauvais, Nicolas Kuszla, Elitsa Ivanova, Marie Simon, Adrien Dufour, Frédéric Launay, Florence Gazeau, Hervé Acloque, Sophie Parat, Joël Poupon, Anne Burtey

## Abstract

The sustainability of mammals on Earth relies on milk. During lactation, maternal exposure to pollutants like metal nanoparticles (NPs) can affect offspring development and survival. Despite being banned from food applications in Europe due suspected toxicity, titanium dioxide (TiO_2_) NPs are still massively manufactured for countless other uses. While contamination of ecosystems is well documented, contamination of mammals remains underexplored. Here, we used synchrotron X-ray fluorescence and single particle inductively coupled plasma mass spectrometry to analyse human, animal, and infant formula milk. Titanium containing micro- and nano-particles were detected in all samples, regardless of the species, location, and processing. We identified varying concentrations, sizes, and combinations of rutile and anatase TiO_2_, ilmenite FeTiO_3_ and possibly titanite CaTiSiO_5_ or pseudobrookite Fe_2_TiO_5_. These findings suggest that milk serves as a carrier for titanium-containing nanomaterials to expose newborns on a daily basis until weaning.

## Introduction

Engineered titanium dioxide nanoparticles (TiO_2_-NPs) are massively used in all industrial sectors ^1,2^. Anatase and rutile TiO_2_-NPs are typically manufactured from ilmenite (FeTiO_3_) ores for their coloring, photocatalytic, UV-filtering, or anti-corrosive properties ^1,3^. They are employed in countless applications ranging from cosmetics, pharmaceutics, electronics, paints to even food ^1,2,4–6^. Use and waste degradation of Ti-containing products along with their release as industrial by-product result in their unintentional release in ecosystems ^2,5,7–13^ while they are intentionally released on crops and soils for their fertilizer properties ^14–16^, added to wastewater as remediation agents ^17,18^ and even proposed for artificial ocean fertilization ^19^. Emergent literature reported the presence of TiO_2_-NPs in ecosystems. They were detected in Austrian lake waters ^9^, Australian and Dutch rivers ^20–22^, sea water in Brazil ^12^ Australia ^21^ and Spain ^10^, atmospheric particulate matters of all sizes in Brazil ^13^ and The Netherlands ^20^, landfill soils in the US ^23^ and Switzerland ^8^, and in sewage sludges and soils amended with the latter in the US ^7,17,20,23,24^. The first evidence for the contamination of living organisms by Ti-NPs has been provided recently in fish ^12,13^ and in human colon tumor sections ^25^, placenta, and newborn first stools ^26^, the latter indicating materno-foetal transfer. Whether such contamination represents a health hazard is unclear. Ti-NP toxicity remains debated and is likely to depend on the mineral phase, size, and concentration of nanoparticles^27–29^. However, TiO_2_-NPs were shown to exert geno- and cyto-toxic effects *in vitro* and *in vivo* ^1,4,28,30–32^, including upon oral exposure in rodents ^31^, and proposed to participate in pathological disorders such as cancer ^25,33–35^, inflammatory gut ^36,37^, neurological and cardiovascular diseases ^38,39^, asthma ^40,41^ and microbiota dysbiosis ^42,43^. They were classified as possible carcinogenic to humans upon inhalation by The International Agency for Research on Cancer (IARC) ^34,44^ and their used in food application as a colorant (E171) has been banned in France in 2020 and in Europe in 2022 ^32^. If their toxicity is debated in adult organisms, little is known about their toxicity in the fragile perinatal period. In *C. elegans*, exposure of newborn worms to TiO_2_-NPs altered their development, locomotion, and reproductive functions ^45^. In young rodents, exposure to TiO_2_-NPs has been shown to induce gut dysbiosis and intestinal immune injuries ^46^. Interestingly, in lactating rodents, maternal exposure altered the development, growth, and survival of their offspring ^47–50^. These health effects were shown to correlate with elevated soluble Ti levels measured in acid-digested milk samples by ICP-MS ^47^, hence without providing evidence for the presence of Ti as Ti-containing NPs (Ti-NPs). To address the existence and the extent of actual contamination of milk by Ti-NPs, we used a combination of state-of-the-art synchrotron-based X-ray spectroscopy and mass spectroscopy approaches. We explored the presence, concentration, speciation (determination of chemical species), and size in the micro- and in nano-range, of Ti in samples of human, dairy livestock, and infant formula milk.

## Results

### Titanium element is detected in milk

First, we used Inductively Coupled Plasma Mass Spectrometry (ICP-MS) to evaluate the presence of Ti in milk. We analyzed infant formula (IF) and dairy livestock milk (Table S1). IF samples were from several manufacturers intended for newborns (0-6 months) or infants (6 to 36 months), thickened or not, organic-based or not (Table S2). Dairy livestock powder milk (PM) was raw or skimmed, organic or not, from cows or goats and commercialized in France, Austria, or Poland (Table S1-2). Given that ICP-MS detects soluble Ti, we mineralized milk samples by double nitric-hydrofluoric acid digestion to release Ti from Ti-containing particles. As shown in Fig. 1A, Ti was detected in all the IF samples tested and in 1 out of 5 PM samples. The concentration of Ti in IF samples ranged from 1.13 to 50.72 µg per liter of milk, with an average of 13.35 +/-3.85 µg Ti/L which was much lower in PM (0.68 +/-0.54 µg/L (Fig.1A)). Of note, Ti concentration was similar in batches of IF acquired several months apart (compare IF1 vs. IF1’, IF5 vs. IF5’, IF6 vs IF6’, Fig. 1A). To verify that the absence of detection of Ti in some samples was not due to a suboptimal mineralization, Ti signal was acquired 1 000 times with a very short reading time of 100 microseconds (µs) instead of 50 ms used in classical analytical conditions. In these conditions, any particles present would appear as elevated transient signals. Complete dissolution of Ti was confirmed in all milk samples, even in those with low Ti concentration. An example is given in Fig. S1 showing a profile of Ti signal similar to that of a standard solution of Ti.

**Figure 1.**
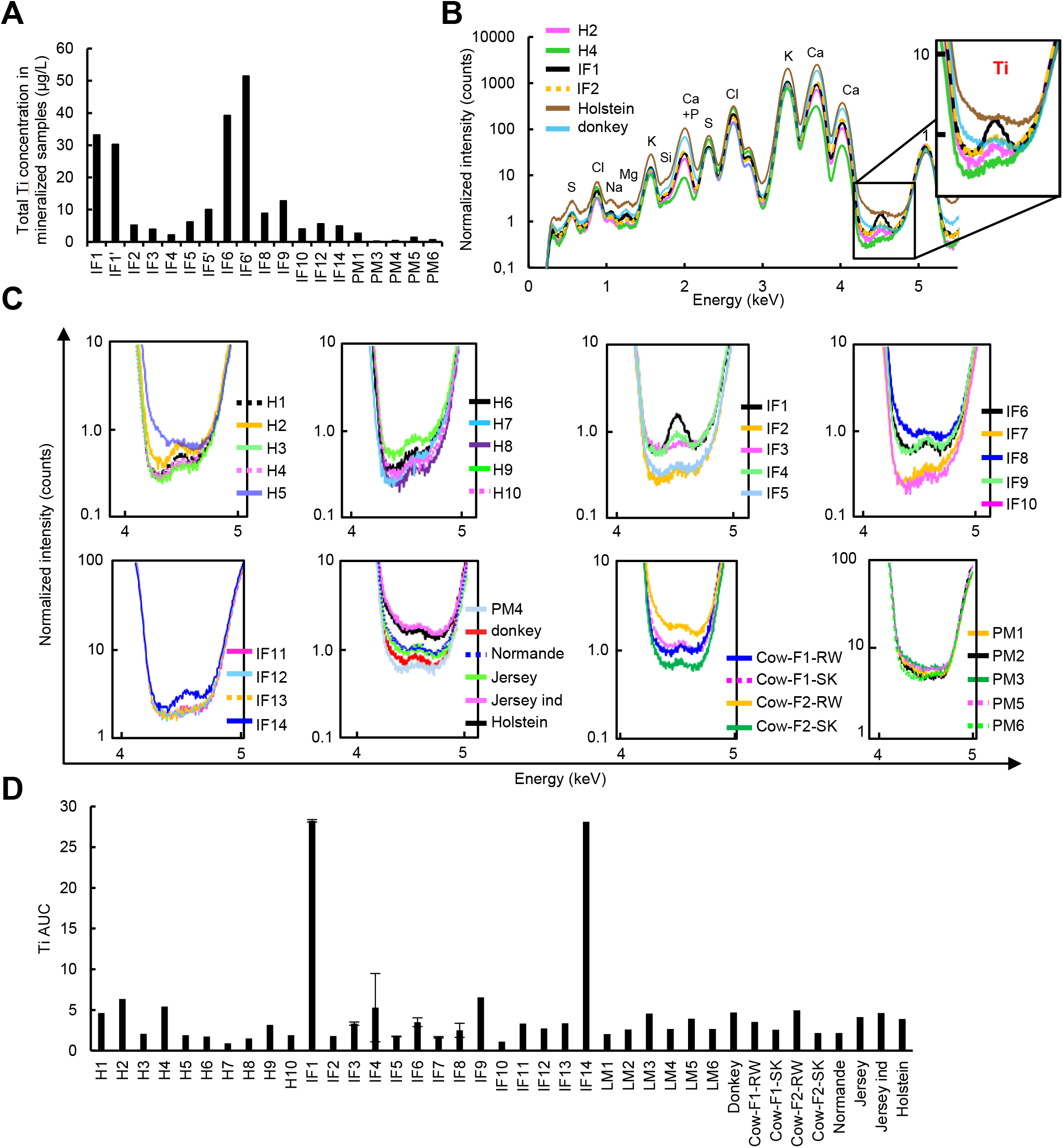
Titanium (Ti) element is present in milk samples. (**A**). ICP-MS measurements of Ti concentration on nitric and hydrofluoric acid digested infant formula (IF) and animal samples including powder milk (PM). (**B**). X-ray fluorescence spectra collected at 5.1 keV on human (H2, H4), IF (IF1, IF2) and animal milk (donkey, Holstein cow milk). Insets are magnifications of Ti peaks. (**C**). Magnifications of Ti peaks for all samples analyzed. (**D**). Area under curve (AUC) of Ti peaks computed by PyMCA software for all samples.

Next, we used synchrotron X-ray fluorescence (XRF) spectroscopy at LUCIA beamline, Synchrotron SOLEIL ^51^ (Saint-Aubin, FR) to investigate the presence of Ti in non-mineralized milk samples. Milk samples that were not already in powder were freeze-dried and pressed into pellets. In addition to the aforementioned milk samples, we analyzed breastmilk samples from ten women living in Paris and its immediate suburbs (collected by the Lactarium Port-Royal, AP-HP, Paris, FR). We also analyzed additional animal milk samples including raw (RW) or skimmed (SK) cow milk from two farms located approximately 20 km from Paris (cow farm 1 “cow-F1” and cow farm 2 “cow-F2”, table S1); organic raw milk from a donkey in Ariège county (FR)(“Donkey”, table S1); raw milk from three breeds of cows (Normande, Jersey, and Prim’Holstein (“Holstein”)) from an INRAE experimental unit (Le Pin-au-Haras, Normandie, FR) located 200 km from Paris, FR (Table S1-2). XRF analysis at 5.1 keV with an unfocused beam revealed the bulk elemental composition of milk samples ^52^ (Fig. 1B and S2). The relative abundance of sulfur (S), chlorine (Cl), sodium (Na), magnesium (Mg), aluminum (Al), potassium (K), silicon (Si), calcium (Ca), phosphorus (P) varied between samples (Fig. 1B-C and S2-3), even when milk originated from the same manufacturer (Fig. S4). Importantly, we observed the presence of a peak at 4.5 keV indicating the presence of Ti (Fig. 1B-C and S2-4) as confirmed by the Ti peak area under curve (Ti AUC, Fig. 1D). Ti was detected in all the samples but with levels varying by a 25-, 7- and 3-fold in IF, human and animal milk respectively (Fig. 1D). Samples with the highest Ti AUC values were IF1, IF14 and IF9, all thickened milk intended for newborns with reflux (Fig. 1D). Amongst the samples with the lowest AUC we found organic-based milk (IF-2, -10, -11, -12, -13, PM3 and donkey), despite the lack of statistic significant difference with non-organic samples (3.08 +/-0.66 vs. 4.76 +/-1.12 in organic vs. non-organic respectively). Even if powder milk were industrially processed, they contained levels of Ti within the same range as those found in farm milk (Fig. 1D). Interestingly, skimmed milk from both farm 1 and 2 cows had lower Ti AUC than their raw counterparts (compare “Raw” vs. “SK” in Fig. 1D) suggesting that Ti may be partly carried by milk fat globules (cream) or somatic cells, both removed by the low-speed centrifugation skimming process. Analysis of milk from cows raised in experimental conditions showed that blended milk from 6 Normande cows contained less Ti compared to blended milk from 6 Holstein or 6 Jersey cows (Fig. 1C and S3F), as confirmed by analyzing Ti AUC (Fig. 1D). Note that milk from a single individual Jersey cow (“ind”, Fig. 1D) had similar Ti AUC level than the blended milk from 6 Jersey cows (Fig. 1D). No clear relationship was observed between Ti levels and milk yield, fat and cells while a slight tendency to correlate was observed between Ti levels and milk protein levels (Fig. S5). Acquisition of several regions of the same milk pellet resulted in similar Ti AUC (IF3: AUC=3.32 +/-0.02, n=4 regions; IF5: AUC=1.75 +/-0.04 n=2 regions; IF7: AUC=1.65 +/-0.09 n=2 regions, Fig. S6A). This homogeneity within samples was however not observed for IF4 (Ti AUC=5.28 +/-4.2, n=2 regions, Fig. S6A), a heterogeneity observed for other elements as well (Fig. S6B). It has recently been suggested that IF may contain silicon dioxide NPs ^53^. Accordingly, we detected the presence of Si in several milk samples, in particular in industrially processed milk samples (PM1-3, PM5 and IF6, IF8, IF9, IF11-14) and to a lesser extent in human and farm milk (Fig. S7).

### Titanium distributes in hotspots by micro-XRF

We used micro-XRF analysis to investigate the presence and distribution of Ti particles in milk pellets. We acquired maps of 1 to 3 mm^2^ in continuous flyscan mode with a step size of 3 µm and analyzed human (n=8), IF (n=4) and livestock (n=3) milk samples (Fig. 2, S8-11). Our results confirmed the presence of Ti in all samples and revealed that high Ti signal accumulated in small regions that we referred to here as “hotspots” (Fig. 2A, open arrowheads and Fig. S8A), emerging from a diffuse and low Ti signal (Fig. 2B, lineascan). Ti distribution in hotspots was observed in all samples analyzed (Fig. S8-11) and did not match that of other elements such as Ca and Si (Fig. 2C). We used a tailored protocol on the open-source ICY-BioImage analysis software ^54^ to automatically detect and quantify Ti hotspots based on gaussian-filtered Ti maps (Fig. 2D-F and S8B). To avoid the quantification of hotspots in which the level of Ti would be too low to emerge from the surrounding low Ti signal, we only quantified hotspots whose average Ti signal intensity was superior by a 1.5 fold to that of the total image (Fig. S8C, pink bars, ratio “R”>1.5). This conservative threshold excluded hotspots with low Ti intensity levels (Fig. S8C, compare R<1.5 vs manual detection or R<1) for which Ti accumulation could not be confirmed on the average pixel extracted XRF spectrum (data not shown). Therefore, this method slightly underestimated the number of Ti hotspots (Fig. S8C). On Ti maps of samples H1, H3 and H9, the majority of hotspots had very low Ti levels that were too close to that of the surrounding diffuse signal, leading to the detection of hotspots whose Ti content could not be confirmed by observation of the pixel-selected XRF spectrum (data not shown). To avoid this, hotspots were instead detected manually on thresholded images by ImageJ program ^55^ (“manual” Fig. S8C and S11). As shown in Fig. 2D, milk were polydispersed in terms of Ti hotspots intensity except for human H2, 4, 8, 9 and 10 whose Ti hotspots were all rather dim. Hotspots sizes ranged from 1 pixel to 227, 118 and 531 pixels in human, IF and livestock milk respectively. Note that hotspots of 1 pixel in size could be equal or smaller than the beam size (3.1×2.6 µm). This 1-pixel sized population represented 5% of total hotspots in farm cow milk up to 22% in human sample (Cow-F1-RW vs. H10, Fig. 2E,), and in average 13 +/-2.8% in human, 10+/-1.3% in IF and 9 +/-3.8 % in animal milk samples. Taken together by type of milk, sizes of hotspots distributed similarly (Fig. 2E, right). Given that freeze-dried and commercial milk were all pressed from 100 mg dried milk into 78.5 mm^2^ pellets, we could estimate the number of hotspots per gram of milk (Fig. 2F). This calculation was done based on the assumption that the beam had penetrated the whole 1 mm thickness of the pellet, which most likely generates an underestimation of the results. We found that human milk contained 10 952 to 160 139 Ti particles/g (H1 and H2 respectively, Fig. 2F), a 15-fold variation that was higher than variations observed in IF and animal milk (3- and 1.9-fold respectively, Fig. 2F). The average number of Ti particles per gram was similar in human and animal milk (6.93 +/- 2.13 × 10^4^ vs. 7.59 +/- 1.37 × 10^4^ part/g respectively) which tended to be higher in IF samples despite the lack of statistical significance (1.70 × 10^5^ +/- 3.86 × 10^4^ part/g in average in IF).

**Figure 2.**
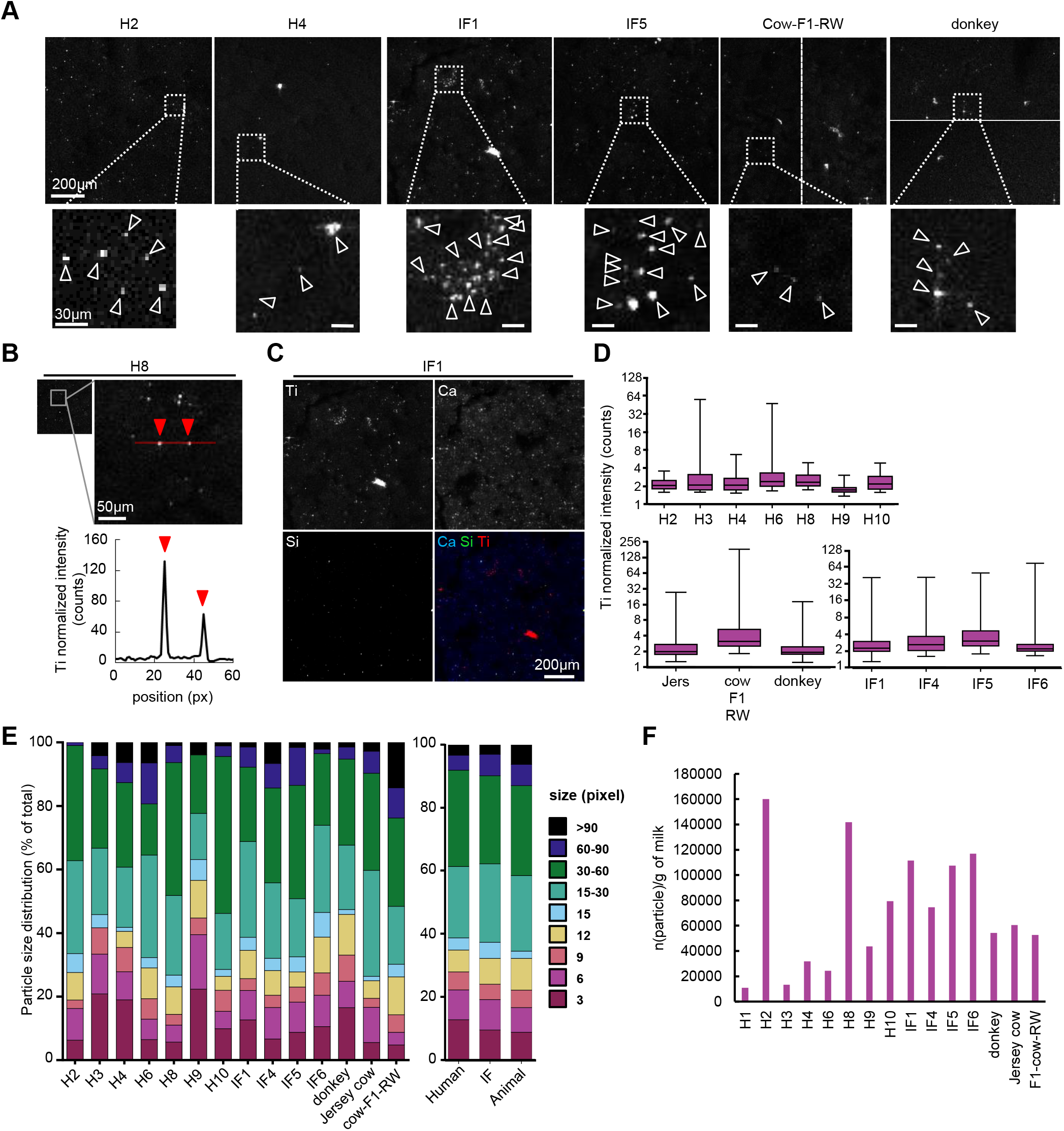
Titanium accumulates in hotspots. (**A**). Micro-XRF Ti maps of human samples (H2, H4), infant formula (IF1, IF5), cow-F1-RW and donkey milk acquired in flyscan mode with a focused 3 × 3μm2 beam. Regions of interest (dotted square) were magnified, open arrowheads point to Ti accumulations in hotspots. Light corresponds to a high Ti level, while dark corresponds to a low Ti level. (**B**). Magnification of a region of interest (dotted box) on H8. Ti intensity is plotted along a line of interest (in red) crossing two hotspots (lower graph). Red arrowheads point to peak of Ti intensity in hotspots. (**C**). Elemental maps of calcium (“Ca”, in blue), silicon (“Si”, in green) and Ti (in red) for human sample H2. (**D-E**). Characterization of hotspots (“particles”) for Ti intensity (D) and size (E). Distribution by size is shown as a % of total hotspots for each sample (E, left graph) or by types of milk (E, right graph). Total area analyzed was as follow: 2.07 mm^2^ for IF4; 2 mm^2^ for IF1, donkey, cow-F1-RW; 1.75 mm^2^ for H8; 1.5 mm^2^ for H2 and 1 mm^2^ for H6, H9, H10, IF5, IF6 and Jersey cow. (**F**). Number of particles per gram of dried milk.

### Chemical state speciation by micro-XANES of hotspots reveals the presence of various titanium-bearing minerals

To speciate the chemical state of Ti in hotspots, we used Ti K-edge µ-XANES ^56^ (Fig. 3 and S12-13). Of the 119 hotspots analyzed, the great majority (94%) resulted in a Ti speciation. Surprinsingly, we detected a diversity of Ti bearing minerals: rutile TiO_2_, anatase TiO_2_, ilmenite Fe_2_TiO_3_ and titanite CaTi(SiO_5_) or pseudobrookite Fe_2_TiO_5_ (“PB”) ^57,58^(Fig. 3A-B and Fig. S12-13). Rutile TiO_2_ accounted for the majority of the Ti minerals detected (65% of total hotspots analyzed) while anatase was the second most represented Ti bearing mineral (17% of total hotspots). Ilmenite represented 5% of total hotspots and PB or titanite in 2%, while the mineral bearing Ti remained not determined for 12% of hotspots analyzed (“nd”) (Fig. 3C). Sample-by-sample analysis showed that each milk samples had a specific combination of Ti bearing minerals. In human milk, there was a majority of rutile TiO_2_ hotspots which were accompanied by a few hotspots of anatase, pseudobrookite or titanite or nd while in some IF samples, only rutile hotspots were detected (Fig. 3C, compare all H with IF4). In another IF sample, we detected a majority of ilmenite hotspots, a few anatase and no rutile (IF6, Fig. 3C). Note that ilmenite was not found in any other milk sample analyzed. Anatase, which represented 17% of all the hotspots in human samples, represented 100% of IF4, 50% of IF1 and 33% of IF6. Regarding farm milk, donkey milk had a mix of *(58:25:17) (*rutile:anatase:nd) and normande cow milk was analyzed only for one hotspot that was rutile TiO_2_ (Fig. 3C and S13). Taken together, the global composition of Ti bearing minerals was as follows: (%rutile:anatase:ilmenite:PB:nd): animal (73:10:0:2:15), human (79:13:0:0:8) and IF (53:25:17:3:2) (Fig. 3C and S12-13).

**Figure 3.**
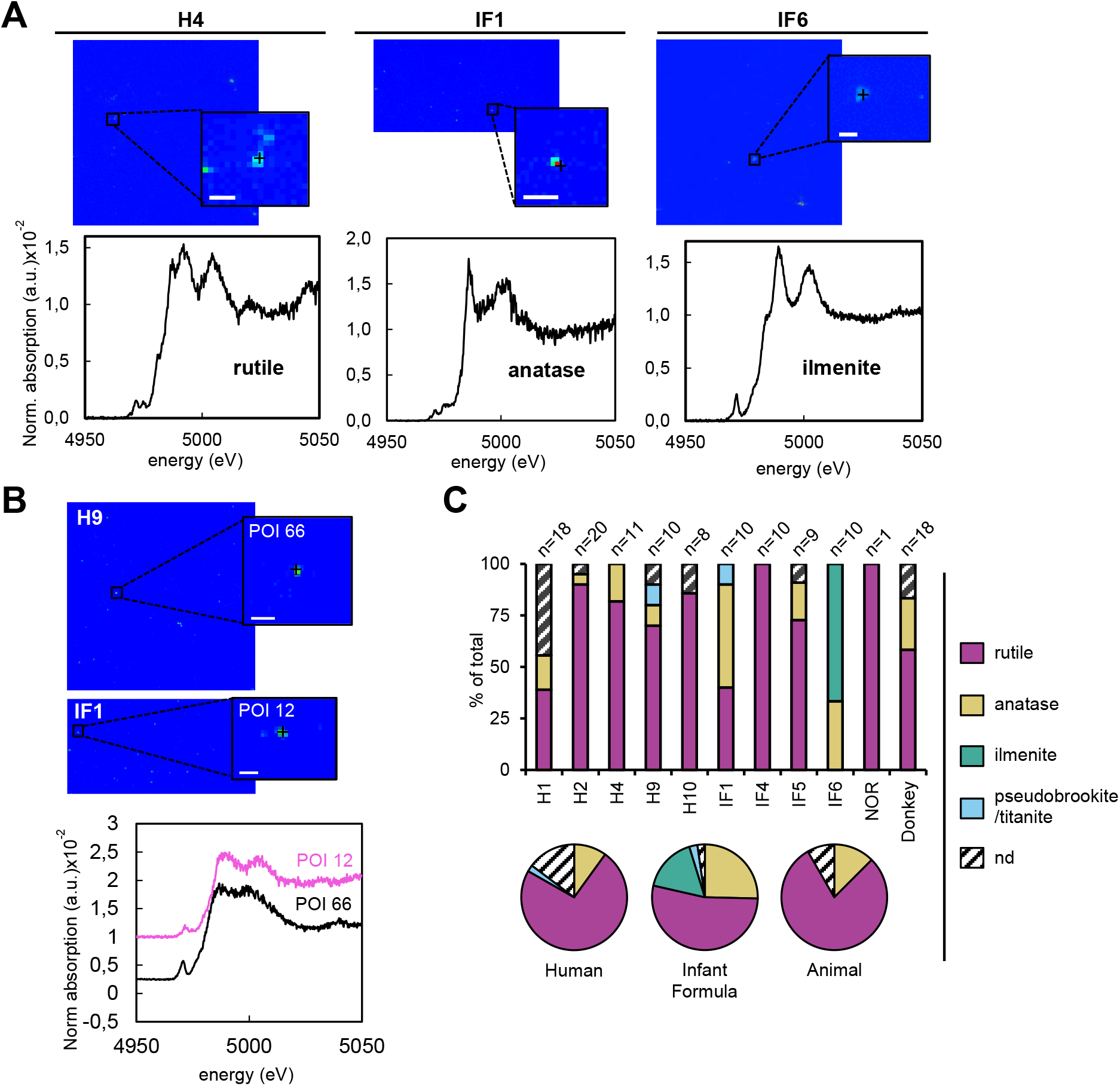
Ti hotspots contain rutile (TiO_2_), anatase (TiO_2_), ilmenite (Fe_2_TiO_3_) and possibly pseudobrookite Fe_2_TiO_5_ or titanite CaTiSiO_5_. (A) Hotspots on H4, IF1 and IF6 were analyzed by Micro-X-ray absorption Near Edge Structure (µXANES) revealing spectra of rutile, anatase and ilmenite. Magnification around a region of interest is shown (black dotted lines) and the POI analyzed is marked by a black cross (B) µXANES spectra of POI 12 and 66 on H9 and IF1 Ti maps. (C) Percentage of Ti mineral phases detected in hotspots for each sample analyzed or as an average by type of milk. Scale bars 25µm.

### Titanium-containing particle sizes are in the nano-range

Next, we investigated the presence of Ti-containing nanoparticles (Ti-NPs) by using the Single Particle ICP-MS (SP-ICP-MS) Nano Application Module of the Syngistix™ software. Milk samples were only diluted 1/10^th^ in ultrapure water. Samples insufficiently dissolved to pass through the instrument capillaries without clugging them were centrifuged at low speed to remove larger clumps (IF1, IF1’, IF5, IF5’, and IF8, Table S3) or heated and sonicated for those that were too grainy (PM1, -4, and -6, Table S3). SP-ICP-MS consisted of 50 µs acquisitions of Ti intensity. By analyzing 1 000 000 measurements, results demonstrated the presence of Ti-containing particles as high Ti intensity peaks that emerged from a lower Ti intensity signal corresponding to Ti ions (“particles” vs. “soluble”, Fig. S14). Ti intensity peaks were converted into TiO_2_ particle diameter by the Syngistix Nanomodule assuming that particles are spherical. This approximation to particle shape is essential for calculating particle size, this calculated diameter will be used in the following results. As shown in Fig. 4A and S15, all milk samples contained nanosized Ti particles. Their average diameter was 151.7 +/-13.3 nm in IF and 132.8 +/-6.9 nm in animal milk (Table S3), with size distributions varying between samples (Fig. S15). Considering that the generally accepted definition of a nanoparticle states that at least one dimension of the particle should be below 100 nm in size ^2^, we analyzed the percentage of particles below 100 nm. In IF, Ti-NPs represented a third of the total Ti particles detected (average 31.9 +/ 7.5 %, Fig. 4B). Some samples contained 99 % of Ti-NPs (IF5) while others hardly any (IF3: 4 %, table S3 and Fig. S15). In dairy livestock PM, Ti-NPs represented 38.3 +/-7.8% of the total Ti particles (Fig. 4B and Table S3). We then analyzed the concentration of Ti particles per gram of milk and liter of milk after resuspension in water following manufacturer recommendations to reconstitute bottle IF and PM milk (Fig. 4C-D). We found that IF contained in average 4.01 +/-2.91 ×10^8^ Ti part/L while dairy livestock milk contained an average of 1.25 +/-0.54 × 10^8^ Ti part/L. IF5 and IF10 had the highest concentration of particles (3.99 × 10^9^ and 4.29 × 10^8^ part/L respectively), the lowest being IF8 and IF1’ (4.83 × 10^6^ and 9.68 × 10^6^ part/L respectively (Fig. 4C-D and Table S3). The two batches of IF5 acquired several months apart showed very different Ti particle levels, while it was similar for IF1 and IF6 batches (IF1 vs. IF1’ compared to IF5 vs. IF5’ and IF6 vs. IF6’, Fig. 4C-D, Table S3). A solution of 10 µg/L TiO_2_-NP standard reference material (Titanium Dioxide Nanomaterial 1898, National Institute of Standards & Technology) ^59^ was used as a positive control. We found it contained 1.47 × 10^8^ part/L with a median particle size of 326 nm and the absence of particles below 100 nm (Table S3) in accordance with the certificate of analysis for a non-sonicated solution^59^. To verify the presence of particles of Ti and dissolved Ti in samples, we hypothesized that particles should be retained upon ultrafiltration on 30 kDa filters while soluble Ti should pass through. Accordingly, the concentration of Ti particles decreased upon filtration (Fig. S16A), in IF5’ it dropped from 5.08 × 10^7^ to 0.48 × 10^7^ part/L and in IF4 from 2.47 × 10^8^ to 0.10 × 10^8^ part/L (Fig. S16 and Table S3). Ultrapure water contained 8.6 × 10^5^ part/L (“ultrapure water”, Fig. 4D, S16B, table S3), a concentration that was increased upon ultrafiltration to 4.6 × 10^6^ part/L indicative of possible contamination during the process (Fig. S16B). To address whether some particles may be aggregates of NPs, we used sonication known to dissociate aggregates of NIST TiO_2_-NPs ^59^. Sonication of the NIST standard induced a decrease of the Ti particle median size from 326 to 186 nm along with an increased concentration from 1.4 × 10^8^ to 7.0 × 10^8^ part/L before and after sonication respectively that corresponds to a fold change of 4.8 (Table S3). Similar trends were observed upon sonication of IF3 and IF5’ whose median particle size decreased from 146 to 96 nm and from 116 to 106 nm respectively while the number of particles increased slightly for IF3 and by 2.5-fold for IF5’ (Table S3). Finally, the contribution of soluble vs. particle Ti to the total Ti concentration varied between samples (Fig. 4E and Table S3).

**Figure 4.**
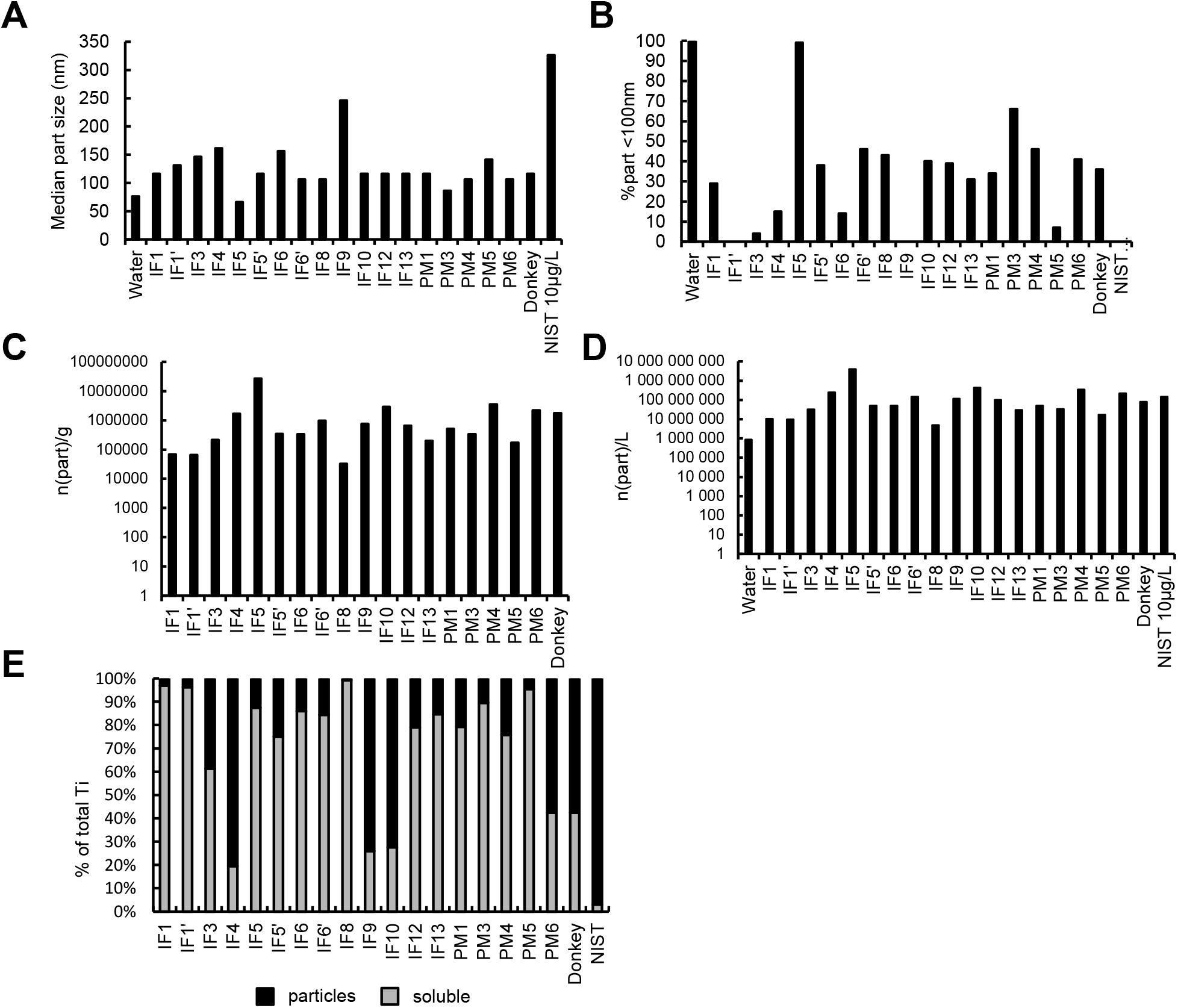
SP-ICP-MS reveals the presence of Ti nanoparticles in milk. IF and animal milk samples were diluted in water (1/10^th^) and analyzed along with ultrapure water and a 10 µg Ti/L solution of standard TiO_2_-NP (NIST) by single nanoparticle ICP-MS. (A) Ti particle size, (B) % of total particles with a size below 100 nm and (C-D) number of Ti particles per gram of milk (C) or per liter of reconstituted milk, water or NIST solution. (E) Participation of soluble and particle to Ti concentration.

## Discussion

As milk is an absolute requirement for mammals, it has been closely monitored for toxic compounds for years. Recently, a study has shown the contamination of breast milk by several toxic chemicals including per- and poly-fluoroalkyl substances (PFASs)^60^ but neither this study nor any of the literature we are aware of ^61–64^, has investigated the presence of titanium element or of Ti-NPs in milk and more generally, the passage of nanomaterials through the mammary gland into milk. Here we have demonstrated the existence of a widespread contamination of milk by soluble titanium and also by titanium-containing microparticles and nanoparticles in human, dairy livestock and industrial milk. We have provided an extensive characterization of its chemical composition and mineral phase, as well as its particle size and concentration. Recently, it has been shown that oral exposure of lactating mice to TiO_2_-NPs resulted in increased levels of Ti in milk ^47^. However, this study did not provide evidence for the presence of Ti-containing particles in milk given that authors mineralized milk samples before ICP-MS analysis ^47^. Here, using µXRF at synchrotron SOLEIL and SP-ICP-MS, we revealed the presence of Ti-containing microparticles and nanoparticles respectively. Whether the former are aggregates of the latter was shown for Ti microparticles collected in rivers and atmospheric matter ^2,11,13,65^ and is supported by the decrease of Ti particle size that we observed upon milk samples sonication. SP-ICP-MS analysis also showed the presence of soluble Ti in milk, and that the contribution of soluble vs. particles to the total Ti concentration varied between samples. Note that the sample introduction system limits the maximal size of particles detected to 10 microns ^66^, thereby not detecting all the microparticles above that size, but these were detected by µXRF.

Ti presence was demonstrated in all analysed human, dairy livestock, and infant formula milk analyzed. Similar Ti levels were measured in donkey milk originating from a farm in Ariège county (South of France) compared to cow milk from farms located 20 km from Paris while the levels varied in breastmilk despite women all lived in the same area (Paris or its immediate suburbs). Larger-scale analysis is therefore needed to investigate whether the contamination of milk by Ti is universal or whether it varies according to the geographical location of the sampling.

The source of Ti in milk remains unclear. TiO_2_-NPs are massively used in all industrial sectors but the use of ilmenite, pseudobrookite, or titanite, which could explain their presence in milk, is less documented. In infant formula, Ti may originate from cow milk but given that the latter had lower Ti levels than IF, other sources of Ti contamination are likely to exist during the industrial processing for example. In dairy livestock, it is tempting to speculate that milk Ti derived from environmental sources in which Ti has been detected (water, soil, and air ^2,5,7,8,11–13,17,20–23^). Additional sources may account for Ti in breastmilk such as daily life products (cosmetics, toothpaste, food, pharmaceuticals)^1,6^, eroding building paints and coats, industrial activity, traffic, or proximity to soils and crops treated with Ti-NPs as fertilizers ^1,5,13–16^. Differential exposure to these sources may explain the observed variability of Ti levels in human milk.

TiO_2_-NPs can be absorbed by different routes ^34,67^. In lactating rodents, both the airway and oral exposure to TiO_2_-NPs increased the concentration of Ti element in milk, but the airway resulted in higher levels in milk ^47^ suggesting that observed Ti levels variations in breastmilk could be the result of differences in exposure routes. Maternal physiological factors and genetics may also influence milk composition and yield ^68^. We observed less Ti in Normande milk compared to the two other breeds despite all cows were raised in identical conditions in an INRAE experimental unit. Normande cow produced less milk than Jersey and Holstein cows arguing against a dilution due to larger volume of milk produced by Normande cows. Mice exposed to TiO_2_-NPs during lactation had elevated soluble Ti element in milk somatic cells ^47^. Here, variations of Ti levels in the three breeds did not seem to correlate neither with that of somatic cells nor with fat content. However removal of cells and fat by low-speed centrifugation of farm milk correlated with a decrease of Ti levels. Additional studies are thereby needed to clarify the role of lipids, cells and proteins as carriers of Ti especially given that Ti levels and milk protein content tended to vary similarly. Caseins, the major family of milk protein are known to assemble into micelles proposed to vehicle minerals and bioactive molecules ^69^. Whether the latter could encapsulate NPs and deliver it to offspring remains unknown. Other mechanisms may also be involved in the secretion of Ti in milk, including a passive transfer across the mammary epithelial barrier, which is damaged upon TiO_2_-NP exposure in rodents ^49,70^. In 2010, TiO_2_-NPs were classified as possibly carcinogenic to humans by inhalation ^34,44^ and banned from use in food in France in 2020 and Europe in 2022 ^32^. Our findings demonstrating the presence of Ti-NPs in milk raise concerns about their impact on neonatal and infant development and health. During lactation, maternal exposure to TiO_2_-NPs provoked multiple organ alterations in neonates breastfed, diminished their memory and learning skills, and decreased their growth and survival rate ^47–49,71^. These *in vivo* studies together with the large number of *in vitro* studies ^4,67^ were based on NPs with different characteristics to those we identified here. Hence, efforts are needed to investigate the effects of Ti-containing particles and NPs with the size distribution, concentration, and chemical composition identified herein in animal and human milk.

## Materials and Methods

### Reagents and consumables

Ultrapure water (resistivity > 18.2MΩ), Milli-Q® Integral3, Merck Millipore, Molsheim, France. Ultrapure nitric acid (67-69% HNO3 PlasmaPURE® Plus ref 250-039-171, SCP Science, Villebon-sur-Yvette). Ultrapure hydrofluoric acid (47-51% HF PlasmaPURE® Plus ref 250-036-121, SCP Science, Villebon-sur-Yvette). Certified Reference Material NCS DC 73347, Human Hair, 2009, China National Analysis Center for Iron and Steel. Standard Reference Material 1898 Titanium Dioxide Nanomaterial, National Institute of Standards & Technology. Centrifugal Filter Units Amicon® Ultra-4® Centrifugal Filters Ultracell 30 K (Merck Millipore, Molsheim, France). Polypropylene high clarity polyethylene (ISO 12485/ISO9001) 15 mL and 50 mL tubes were used to sample IF and animal milk while human milk samples were stored in hospital bottles CE0459 certified. Ilmenite FeTiO3 was purchased from Sigma-Aldrich (cat. NO. 400874).

### Milk samples

Samples of milk analyzed are listed in Tables S1-2. Human milk samples were collected by the Lactarium Port-Royal (AP-HP, Paris, FR) from 10 women living in theParis area. The donkey milk sample was obtained from an organic farm in Ariège county (FR), and cow milk samples were purchased from two farms (“cow-F1”, and “cow-F2”; located approx. 20 km from Paris (FR)). Milk samples from cows were also collected at the INRAE Experimental Unit Le Pin (Normandie region, (FR)). All cows were fed outdoors on fields of ray-grass Lolium perenne and white clover Trifolium repens grass and supplemented by 400 kg/year of OQUALIM-RCNA certified food concentrate consisting of rapeseed and sunflower meal feed, maize, wheat, wheat bran, beet molasses, calcium carbonate, sodium chloride. Milk samples were collected individually at the morning milking on 18/03/2021 from Jersey cows (n=5) (“Jersey”), Normande cows (n=6, “Normande”), or Prim’Holstein cows (“Holstein”). Powder milk (PM) was purchased from grocery stores or online and included organic or not, raw or skimmed cow milk (two brands, (FR)), goat milk (Poland), raw cow milk (Austria) (Table S1). Infant formula (IF) milk was purchased from pharmacies or grocery stores, included first-age (0-6 months) organic or not, second-age (6-12 months old) and infants (6-36 months), thickened or not, organic or not from six manufacturers (Table S2).

### Milk processing and quality analysis

Human milk samples were frozen within 24 h after collection. Raw farm milk samples were centrifuged at 2000 rpm at 4°C for 30 minutes to remove fat and cells, before storage at -80°C (Table S1). Milk samples were handled with particular care to avoid contamination by laboratory equipment. Manipulators avoided using Ti containing cosmetics. Cow milk samples from INRAE UEP were kept refrigerated for 24 h prior to blending and storage at -80°C. A sample from an individual Jersey cow was kept for analysis. Cow milk samples from INRAE UEP were analyzed by infra-red spectroscopy for the levels of proteins and fat and somatic cells were counted according to NF EN ISO 1333662 standard.

### ICP-MS analysis

#### Milk samples preparation for ICP-MS

##### Mineralized milk

200 mg IF, PM, and freeze-dried donkey milk were incubated for 120 minutes at 95°C with 1mL of ultrapure nitric acid and 1mL of ultrapure hydrofluoric acid. After cooling, 8mL of ultrapure water was added.

##### Reconstituted milk

1 g IF, PM, and donkey freeze-dried milk was dissolved in 14mL of ultrapure water and vortexed prior analysis. IF-1, -1’, -5, -5’, and -8 were centrifuged for 10 minutes at 3500 rpm to remove large clumps. PM1, 4, and 6 samples were sonicated for 10 minutes in a water bath and then heated at 70°C for 90 minutes.

#### Total Ti concentration

Titanium was measured by triple quad ICP-MS NexION® 5000 (Perkin-Elmer, Villebon-sur-Yvette, FR) equipped with a Dynamic Reaction Cell (DRC®). As the major isotope of Ti (48Ti) interfered with 48Ca and several polyatomic species^74^, Ti was measured in the MS/MS mode using ammoniac (0,8 mL/min) as reaction gas to form a TiNH(NH_3_)_4_ complex read at mass 131^75^. The first quadrupole was then set at mass 48 and the second quadrupole at mass 131. For sample introduction, we used a PFA MicroFlow nebulizer coupled to a 50 mL baffled cyclonic spray chamber. Acquisition parameters were: Dwell time 50 ms, 20 sweeps, 10 readings, 3 replicates (10 sec integration time per replicate). Mineralized samples were diluted 1:10 in ultrapure water before analysis in duplicates. Accuracy was checked by analysis of a Certified Reference Material (NCS DC 73347) prepared as milk samples: certified value 2.7 ± 0.6 µg/g, found to be 3.3 µg/g.

#### Ti particle

Ti determination using very short dwell time (100 µs): Analysis of reconstituted milk samples with reduced dwell time from 50 ms to 100 µs allowed us to observe some elevated transient signals, identified as Ti-containing particles, emerging from a continued signal of dissolved Ti. Such transient signals were no more observed in mineralized milk indicating a total dissolution of Ti-containing particles.

#### Single particle ICP-MS (SP-ICP-MS)

Measurements were done with a NexION 5000 ICP-MS in single particle mode with the Syngistix™ Nano Application software module. Samples were then analyzed along with a Ti calibration standard with 50 µsec integration time repeated 1 000 000 times. Such a short integration time provides several measurements for a single particle (Fig S13B). Particle size, distribution, and concentration were automatically computed by the software taking into account the nebulization flow rate, the transport efficiency factor, and the approximation of spherical particles.

### X-ray fluorescence (XRF) and micro-X-ray absorption spectroscopy (µXAS)

Frozen milk samples were freeze-dried using a Cosmos -80°C freeze-drier (Cryotec, Saint-Gely-du-Fesc, FR), and mixed to homogenize. 100 mg of milk were pressed into 10 mm diameter pellets that were placed on a copper holder for analysis. The rutile and anatase TiO2 reference spectra were from^58^. Ilmenite was mixed with cellulose to generate an optimal concentration for transmission analyses. XRF and µXANES were performed at the LUCIA (Line Used for Characterization by Imaging and Absorption) beamline^51^ at the SOLEIL Synchrotron radiation facility (St-Aubin, FR). The Si (111) double-crystal monochromator was calibrated by setting the maximum of the first derivative of a metallic Ti foil XANES spectrum at 4.966 keV. An unfocused beam (about 2.6 × 2.1 mm^2^) was used for XRF spectra collection, and the beam was focused to 3.1 × 2.6 µm^2^ (H x V) using Kirkpatrik-Baez mirrors for micro-XRF (µXRF) and µXANES acquisition. The fluorescence signal was collected using a 60 mm^2^ mono-element Si drift diode (Bruker AXS, Madison, USA). The experimental chamber was under vacuum to reduce the absorption and scattering of X-rays through the air.

### Data acquisition and processing

XRF spectra were collected at 5.1 keV in unfocused mode with a total acquisition time of 4 min. The µXRF maps were collected at 5.1 keV over areas of 1000 × 1000 µm^2^ in continuous acquisition (Flyscan) mode with an equivalent step size of 3 µm and an acquisition time of 50 ms/pixel. The fluorescence spectra, collected from each pixel of the map were normalized by the incident flux and corrected from the detector deadtime value using scripts in Jupyter notebook. Elemental maps are then extracted using fast XRF linear fit PYMCA software plugin (50), implemented in a Jupyter notebook, that fitted the XRF spectra in each pixel of the map.

Ti hotspots detection and fluorescence intensity were done automatically based on the open source ICY Bioimage Analysis protocol (52) consisting of Gaussian filtering of images subjected to the wavelet spot detector plugin (Source code: https://gitlab.pasteur.fr/bia/spot-detector-block; Fig S7C et S8-9). Hotspots considered positive were those whose average intensity was superior to that of the average intensity per pixel of the entire image. To that end, the average intensity per pixel of hotspots was divided with that of the image. Hotspots with a ratio above 1.5 were considered positive since using a threshold of 1.0 resulted in hotspots considered positive despite the Ti signal being too close to the background and that could not be confirmed on the pixel-selected XRF spectrum (Fig. S7C). For six maps in total (H9_33680, H1_33518, H1_33530, H1_33588, H1_33602, and H3_33794), hotspot detection was done by manual detection with the open-source ImageJ (53) using the wand tool on thresholded images. The threshold applied was only set for the minimal value, the maximal value being the maximal levels of the image (Fig. S10).

For µXANES analysis, the points of interest (“POI”) were chosen based on µXRF Ti fitted maps. The XANES spectra were recorded in fluorescence and in continuous acquisition (FlyScan) mode over the energy range of 4.95 to 5.05 keV with an equivalent energy step of 0.2 eV and an acquisition time of 200 or 400 ms/energy step, for XANES (ilmenite reference) or µ-XANES respectively. If the signal-to-noise ratio is sufficient to determine the crystal phase by comparing it with the reference spectra, only one spectrum is collected. If the spectrum is too noisy for attribution, a second spectrum is recorded and averaged with the first one. Spectra were processed according to standard procedures using Fastosh software (54). The E0 edge energy was chosen as the maximum of the first derivative of the XANES spectrum. The background was subtracted from the spectra using linear regression in the region before the edge and polynomial regression after the edge to obtain a horizontal signal.

### Statistical analysis

If not indicated otherwise, values are given as average +/-sem.

## Acknowledgments

We acknowledge SOLEIL for the provision of synchrotron radiation on the LUCIA beamline (proposals 99210164 and 20211262). We would like to thank Olga Roudenko from the GRADES group at SOLEIL for her help in normalizing and correcting XRF maps. We are grateful to Nathalie Jouan and Sandrine Berton for their work in collecting and providing human breastmilk samples as well as to Frederic Launay at the INRAE experimental unit Le-Pin-au-Haras for providing us with cow milk samples.

## Author contributions

Conceptualization: A.B.

XRF and XANES acquisitions: C.R., A.B., C.H.B, M.S., N.K., H.A., F.G., E.I., A.D.

ICP-MS acquisitions: J.P., N.D.O., R.C, A.B.

Milk collection: S.P. (breastmilk, Lactarium), F.L. (cow milk)

Funding acquisition: A.B., F.G., J.P., C.R.

Project administration: A.B.

Supervision: C.R. (XRF/XANES), J.P. (ICP-MS), A.B.

Writing – original draft: A.B.

Writing – review & editing: A.B., J.P., C.R., F.G.

## Data and materials availability

All data are shown in the main text or the supplementary materials.

## References

1. Buzea, C., Pacheco, I. I. & Robbie, K. Nanomaterials and nanoparticles: sources and toxicity. Biointerphases 2, MR17–71 (2007).

2. Hochella, M. F. et al. Natural, incidental, and engineered nanomaterials and their impacts on the Earth system. Science 363, eaau8299 (2019).

3. Current intelligence bulletin 63: occupational exposure to titanium dioxide. (2023) doi:10.26616/NIOSHPUB2011160.

4. Jovanović, B. Critical review of public health regulations of titanium dioxide, a human food additive. Integr. Environ. Assess. Manag. 11, 10–20 (2015).

5. Baalousha, M. et al. Outdoor urban nanomaterials: The emergence of a new, integrated, and critical field of study. Sci. Total Environ. 557–558, 740–753 (2016).

6. Weir, A., Westerhoff, P., Fabricius, L., Hristovski, K. & von Goetz, N. Titanium dioxide nanoparticles in food and personal care products. Environ. Sci. Technol. 46, 2242–2250 (2012).

7. Laborda, F. et al. Detection, characterization and quantification of inorganic engineered nanomaterials: A review of techniques and methodological approaches for the analysis of complex samples. Anal. Chim. Acta 904, 10–32 (2016).

8. Mitrano, D. M., Mehrabi, K., Dasilva, Y. A. R. & Nowack, B. Mobility of metallic (nano)particles in leachates from landfills containing waste incineration residues. Environ. Sci. Nano 4, 480–492 (2017).

9. Gondikas, A. P. et al. Release of TiO2 Nanoparticles from Sunscreens into Surface Waters: A One-Year Survey at the Old Danube Recreational Lake. Environ. Sci. Technol. 48, 5415–5422 (2014).

10. Tovar-Sánchez, A. et al. Sunscreen Products as Emerging Pollutants to Coastal Waters. PLoS ONE 8, e65451 (2013).

11. Slomberg, D. L. et al. Anthropogenic Release and Distribution of Titanium Dioxide Particles in a River Downstream of a Nanomaterial Manufacturer Industrial Site. Front. Environ. Sci. 8, (2020).

12. Souza, I. da C. et al. Nanoparticle transport and sequestration: Intracellular titanium dioxide nanoparticles in a neotropical fish. Sci. Total Environ. 658, 798–808 (2019).

13. Souza, I. da C. et al. Atmospheric particulate matter from an industrial area as a source of metal nanoparticle contamination in aquatic ecosystems. Sci. Total Environ. 753, 141976 (2021).

14. Cásarez-Santiago, R. G. et al./person-group>. Nanoagriculture and Energy Advances. in Plant Nanobionics: Volume 1, Advances in the Understanding of Nanomaterials Research and Applications (ed. Prasad, R.) 141–164 (Springer International Publishing, Cham, 2019). doi:10.1007/978-3-030-12496-0_7.

15. Sharma, B., Tiwari, S., Kumawat, K. C. & Cardinale, M. Nano-biofertilizers as bio-emerging strategies for sustainable agriculture development: Potentiality and their limitations. Sci. Total Environ. 860, 160476 (2023).

16. Lowry, G. V., Avellan, A. & Gilbertson, L. M. Opportunities and challenges for nanotechnology in the agri-tech revolution. Nat. Nanotechnol. 14, 517–522 (2019).

17. Brar, S. K., Verma, M., Tyagi, R. D. & Surampalli, R. Y. Engineered nanoparticles in wastewater and wastewater sludge – Evidence and impacts. Waste Manag. 30, 504–520 (2010).

18. Puri, N. & Gupta, A. Water remediation using titanium and zinc oxide nanomaterials through disinfection and photo catalysis process: A review. Environ. Res. 227, 115786 (2023).

19. Babakhani, P. et al. Potential use of engineered nanoparticles in ocean fertilization for large-scale atmospheric carbon dioxide removal. Nat. Nanotechnol. 17, 1342–1351 (2022).

20. Bäuerlein, P. S. et al. Is there evidence for man-made nanoparticles in the Dutch environment? Sci. Total Environ. 576, 273–283 (2017).

21. Gonzalez de Vega, R. et al. Analysis of Ti- and Pb-based particles in the aqueous environment of Melbourne (Australia) via single particle ICP-MS. Anal. Bioanal. Chem. 414, 5671–5681 (2022).

22. Peters, R. J. B. et al. Detection of nanoparticles in Dutch surface waters. Sci. Total Environ. 621, 210–218 (2018).

23. Kim, B., Murayama, M., Colman, B. P. & Hochella, M. F. Characterization and environmental implications of nano- and larger TiO2 particles in sewage sludge, and soils amended with sewage sludge. J. Environ. Monit. 14, 1128–1136 (2012).

24. Westerhoff, P., Song, G., Hristovski, K. & Kiser, M. A. Occurrence and removal of titanium at full scale wastewater treatment plants: implications for TiO2 nanomaterials. J. Environ. Monit. JEM 13, 1195–1203 (2011).

25. Gatti, A. M. Biocompatibility of micro- and nano-particles in the colon. Part II. Biomaterials 25, 385–392 (2004).

26. Guillard, A. et al. Basal Ti level in the human placenta and meconium and evidence of a materno-foetal transfer of food-grade TiO2 nanoparticles in an ex vivo placental perfusion model. Part. Fibre Toxicol. 17, 51 (2020).

27. OECD. Titanium Dioxide: Summary of the Dossier. in vol. NO.73 (OECD, Paris, France, 2016).

28. Hou, J. et al. Toxicity and mechanisms of action of titanium dioxide nanoparticles in living organisms. J. Environ. Sci. 75, 40–53 (2019).

29. Shi, H., Magaye, R., Castranova, V. & Zhao, J. Titanium dioxide nanoparticles: a review of current toxicological data. Part. Fibre Toxicol. 10, 15 (2013).

30. Gea, M. et al. Shape-engineered titanium dioxide nanoparticles (TiO2-NPs): cytotoxicity and genotoxicity in bronchial epithelial cells. Food Chem. Toxicol. 127, 89–100 (2019).

31. Trouiller, B., Reliene, R., Westbrook, A., Solaimani, P. & Schiestl, R. H. Titanium dioxide nanoparticles induce DNA damage and genetic instability in vivo in mice. Cancer Res. 69, 8784– 8789 (2009).

32. EFSA Panel on Food Additives and Flavourings (FAF) et al. Safety assessment of titanium dioxide (E171) as a food additive. EFSA J. 19, e06585 (2021).

33. Wang, T. et al. Co-exposure to iron, copper, zinc, selenium and titanium is associated with the prevention of gastric precancerous lesions. Biometals Int. J. Role Met. Ions Biol. Biochem. Med. 36, 1141–1156 (2023).

34. Iarc, W. G. on the E. of C. R. to H. Carbon Black, Titanium Dioxide, and Talc. vol. 93 (World Health Organization International Agency For Research On Cancer, Lyon, France, 2010).

35. Urrutia-Ortega, I. M. et al. Food-grade titanium dioxide exposure exacerbates tumor formation in colitis associated cancer model. Food Chem. Toxicol. 93, 20–31 (2016).

36. Barreau, F., Tisseyre, C., Ménard, S., Ferrand, A. & Carriere, M. Titanium dioxide particles from the diet: involvement in the genesis of inflammatory bowel diseases and colorectal cancer. Part. Fibre Toxicol. 18, 26 (2021).

37. Bettini, S. et al. Food-grade TiO2 impairs intestinal and systemic immune homeostasis, initiates preneoplastic lesions and promotes aberrant crypt development in the rat colon. Sci. Rep. 7, 40373 (2017).

38. Herrera-Rodríguez, M. A. et al. Food-grade titanium dioxide and zinc oxide nanoparticles induce toxicity and cardiac damage after oral exposure in rats. Part. Fibre Toxicol. 20, 43 (2023).

39. Baranowska-Wójcik, E., Szwajgier, D., Oleszczuk, P. & Winiarska-Mieczan, A. Effects of Titanium Dioxide Nanoparticles Exposure on Human Health-a Review. Biol. Trace Elem. Res. 193, 118–129 (2020).

40. Lim, J.-O. et al. Titanium Dioxide Nanoparticles Exacerbate Allergic Airway Inflammation via TXNIP Upregulation in a Mouse Model of Asthma. Int. J. Mol. Sci. 22, 9924 (2021).

41. Deng, R., Zhu, Y., Wu, X. & Wang, M. Toxicity and Mechanisms of Engineered Nanoparticles in Animals with Established Allergic Asthma. Int. J. Nanomedicine 18, 3489–3508 (2023).

42. Radziwill-Bienkowska, J. M. et al. Toxicity of Food-Grade TiO2 to Commensal Intestinal and Transient Food-Borne Bacteria: New Insights Using Nano-SIMS and Synchrotron UV Fluorescence Imaging. Front. Microbiol. 9, (2018).

43. Rinninella, E. et al. Impact of Food Additive Titanium Dioxide on Gut Microbiota Composition, Microbiota-Associated Functions, and Gut Barrier: A Systematic Review of In Vivo Animal Studies. Int. J. Environ. Res. Public. Health 18, 2008 (2021).

44. Baan, R. A. Carcinogenic hazards from inhaled carbon black, titanium dioxide, and talc not containing asbestos or asbestiform fibers: recent evaluations by an IARC Monographs Working Group. Inhal. Toxicol. 19 Suppl 1, 213–228 (2007).

45. Huang, S.-T. et al. Titanium Dioxide (TiO2) Nanoparticle Toxicity in a Caenorhabditis elegans Model. Toxics 11, 989 (2023).

46. Yan, J. et al. Intestinal toxicity of micro- and nano-particles of foodborne titanium dioxide in juvenile mice: Disorders of gut microbiota–host co-metabolites and intestinal barrier damage. Sci. Total Environ. 821, 153279 (2022).

47. Cai, J., Zang, X., Wu, Z., Liu, J. & Wang, D. Translocation of transition metal oxide nanoparticles to breast milk and offspring: The necessity of bridging mother-offspring-integration toxicological assessments. Environ. Int. 133, 105153 (2019).

48. Mohammadipour, A. et al. The effects of exposure to titanium dioxide nanoparticles during lactation period on learning and memory of rat offspring. Toxicol. Ind. Health 32, 221–228 (2016).

49. Zhang, C. et al. Induction of Size-Dependent Breakdown of Blood-Milk Barrier in Lactating Mice by TiO2 Nanoparticles. PLOS ONE 10, e0122591 (2015).

50. Carlé, C. et al. Perinatal foodborne titanium dioxide exposure-mediated dysbiosis predisposes mice to develop colitis through life. Part. Fibre Toxicol. 20, 45 (2023).

51. Vantelon, D. et al. The LUCIA beamline at SOLEIL. J. Synchrotron Radiat. 23, 635–640 (2016).

52. Solé, V. A., Papillon, E., Cotte, M., Walter, Ph. & Susini, J. A multiplatform code for the analysis of energy-dispersive X-ray fluorescence spectra. Spectrochim. Acta Part B At. Spectrosc. 62, 63–68 (2007).

53. VeilleNanos - Sur 23 produits du quotidien testés par AVICENN, 20 contiennent des nanos, non étiquetés, parfois même non autorisés. https://veillenanos.fr/enquete-avicenn-tests-nano-2022/.

54. de Chaumont, F. et al. Icy: an open bioimage informatics platform for extended reproducible research. Nat. Methods 9, 690–696 (2012).

55. Schneider, C. A., Rasband, W. S. & Eliceiri, K. W. NIH Image to ImageJ: 25 years of image analysis | Nature Methods. 671–675 (2012).

56. Landrot, G. FASTOSH: A Software to Process XAFS Data for Geochemical & Environmental Applications. in vol. 1402 (Boston, 2018).

57. Leitzke, F. P. et al. Ti K-edge XANES study on the coordination number and oxidation state of Titanium in pyroxene, olivine, armalcolite, ilmenite, and silicate glass during mare basalt petrogenesis. Contrib. Mineral. Petrol. 173, 103 (2018).

58. Dudefoi, W. et al. In vitro digestion of food grade TiO2 (E171) and TiO2 nanoparticles: physicochemical characterization and impact on the activity of digestive enzymes. Food Funct. 12, 5975–5988 (2021).

59. Holbrook, R. D. SRM No. 1898; Titanium Dioxide Nanomaterial; National Institute of Standards and Technology. (2020).

60. Rovira, J. et al. Mixture of environmental pollutants in breast milk from a Spanish cohort of nursing mothers. Environ. Int. 166, 107375 (2022).

61. Martín-Carrasco, I., Carbonero-Aguilar, P., Dahiri, B., Moreno, I. M. & Hinojosa, M. Comparison between pollutants found in breast milk and infant formula in the last decade: A review. Sci. Total Environ. 162461 (2023) doi:10.1016/j.scitotenv.2023.162461.

62. Boudebbouz, A. et al. Heavy metals levels in raw cow milk and health risk assessment across the globe: A systematic review. Sci. Total Environ. 751, 141830 (2021).

63. Almeida, A. A., Lopes, C. M. P. V., Silva, A. M. S. & Barrado, E. Trace elements in human milk: Correlation with blood levels, inter-element correlations and changes in concentration during the first month of lactation. J. Trace Elem. Med. Biol. 22, 196–205 (2008).

64. Khan, N. et al. Analysis of minor and trace elements in milk and yogurts by inductively coupled plasma-mass spectrometry (ICP-MS). Food Chem. 147, 220–224 (2014).

65. Adam, V. et al. Aggregation behaviour of TiO2 nanoparticles in natural river water. J. Nanoparticle Res. 18, 13 (2016).

66. Matusiewicz, H. Evaluation of Various Types of Micronebulizers and Spray Chamber Configurations for Microsamples Analysis by Microwave Induced Plasma Optical Emission Spectrometry. Chem. Anal.

67. Shi, H., Magaye, R., Castranova, V. & Zhao, J. Titanium dioxide nanoparticles: a review of current toxicological data. Part. Fibre Toxicol. 10, 15 (2013).

68. Golan, Y. & Assaraf, Y. G. Genetic and Physiological Factors Affecting Human Milk Production and Composition. Nutrients 12, 1500 (2020).

69. Tavares, G. M., Croguennec, T., Carvalho, A. F. & Bouhallab, S. Milk proteins as encapsulation devices and delivery vehicles: Applications and trends. Trends Food Sci. Technol. 37, 5–20 (2014).

70. Peng, F. et al. Nanoparticles promote in vivo breast cancer cell intravasation and extravasation by inducing endothelial leakiness. Nat. Nanotechnol. 14, 279–286 (2019).

71. Ebrahimzadeh Bideskan, A. et al. Maternal exposure to titanium dioxide nanoparticles during pregnancy and lactation alters offspring hippocampal mRNA BAX and Bcl-2 levels, induces apoptosis and decreases neurogenesis. Exp. Toxicol. Pathol. 69, 329–337 (2017).

72. May, T. W. & Wiedmeyer, R. H. A table of polyatomic interferences in ICP-MS. At. Spectrosc. 19, 150–155 (1998).

73. Suárez-Oubiña, C., Herbello-Hermelo, P., Bermejo-Barrera, P. & Moreda-Piñeiro, A. Exploiting dynamic reaction cell technology for removal of spectral interferences in the assessment of Ag, Cu, Ti, and Zn by inductively coupled plasma mass spectrometry. Spectrochim. Acta Part B At. Spectrosc. 187, 106330 (2022).

74. May, T. W. & Wiedmeyer, R. H. A table of polyatomic interferences in ICP-MS. At. Spectrosc. 19, 150–155 (1998).

75. Suárez-Oubiña, C., Herbello-Hermelo, P., Bermejo-Barrera, P. & Moreda-Piñeiro, A. Exploiting dynamic reaction cell technology for removal of spectral interferences in the assessment of Ag, Cu, Ti, and Zn by inductively coupled plasma mass spectrometry. Spectrochim. Acta Part B At. Spectrosc. 187, 106330 (2022).

