## Supplementary figures for "Detection of titanium nanoparticles in human, animal and infant formula milk"

**The PDF file includes:**

Figs. S1 to S16
Tables S1 to S3

**Fig. S1.** Acquisition of Ti signal (m/z = 131) of IF1 mineralized milk sample diluted 1:10 by ICP-MS using a dwell time of 100 µs in comparison with a 5 µg/L Ti standard solution. Ti concentration of IF1 measured was 4.42 µg/L.

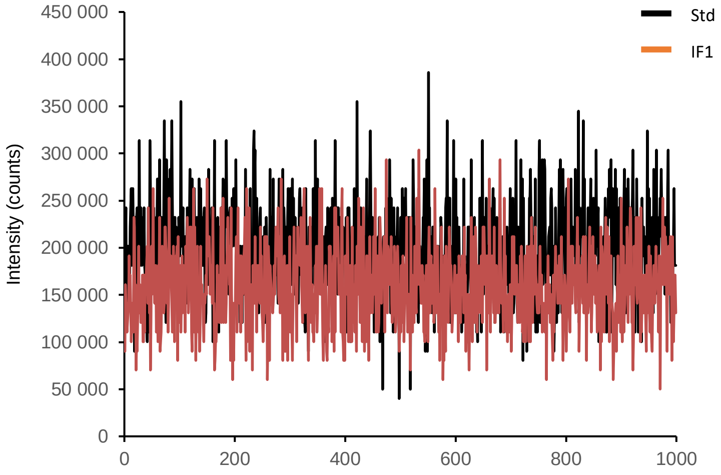

**Fig. S2.** X-ray fluorescence spectrum collected at 5.1 keV on infant formula sample IF6 and corresponding fit using PyMCA software.

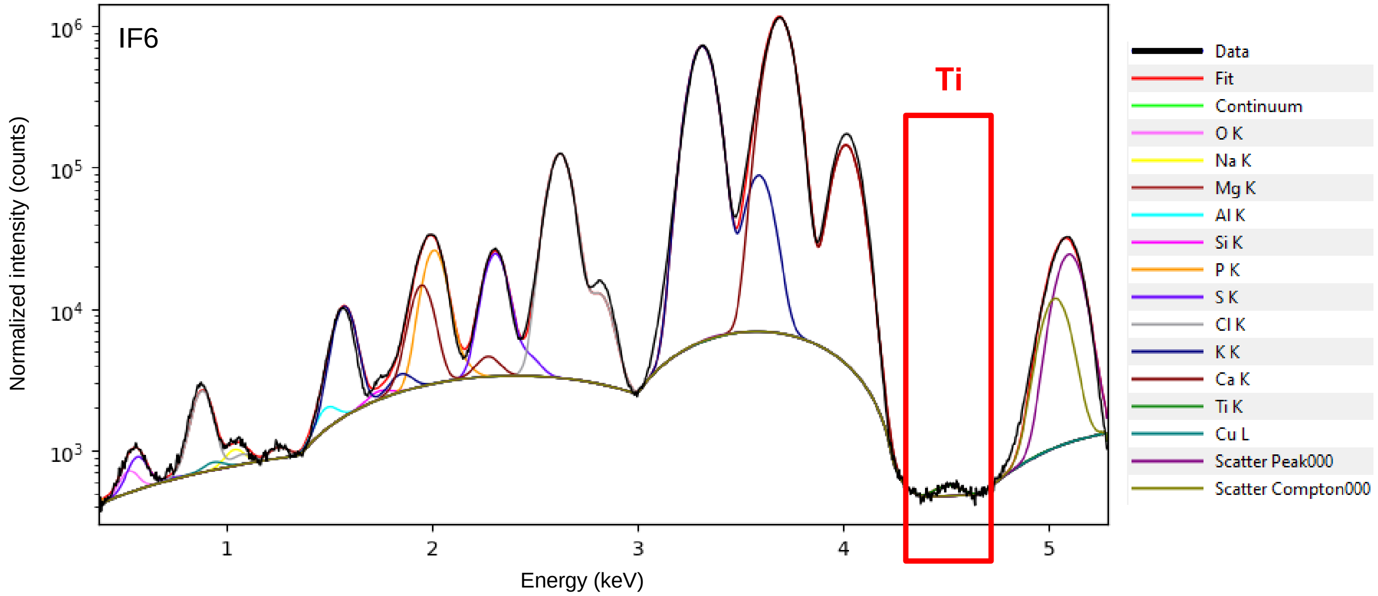

**Fig. S3.** X-ray fluorescence spectra at 5.1 keV of human (A,B), IF (C-E), and animal milk (F-H).

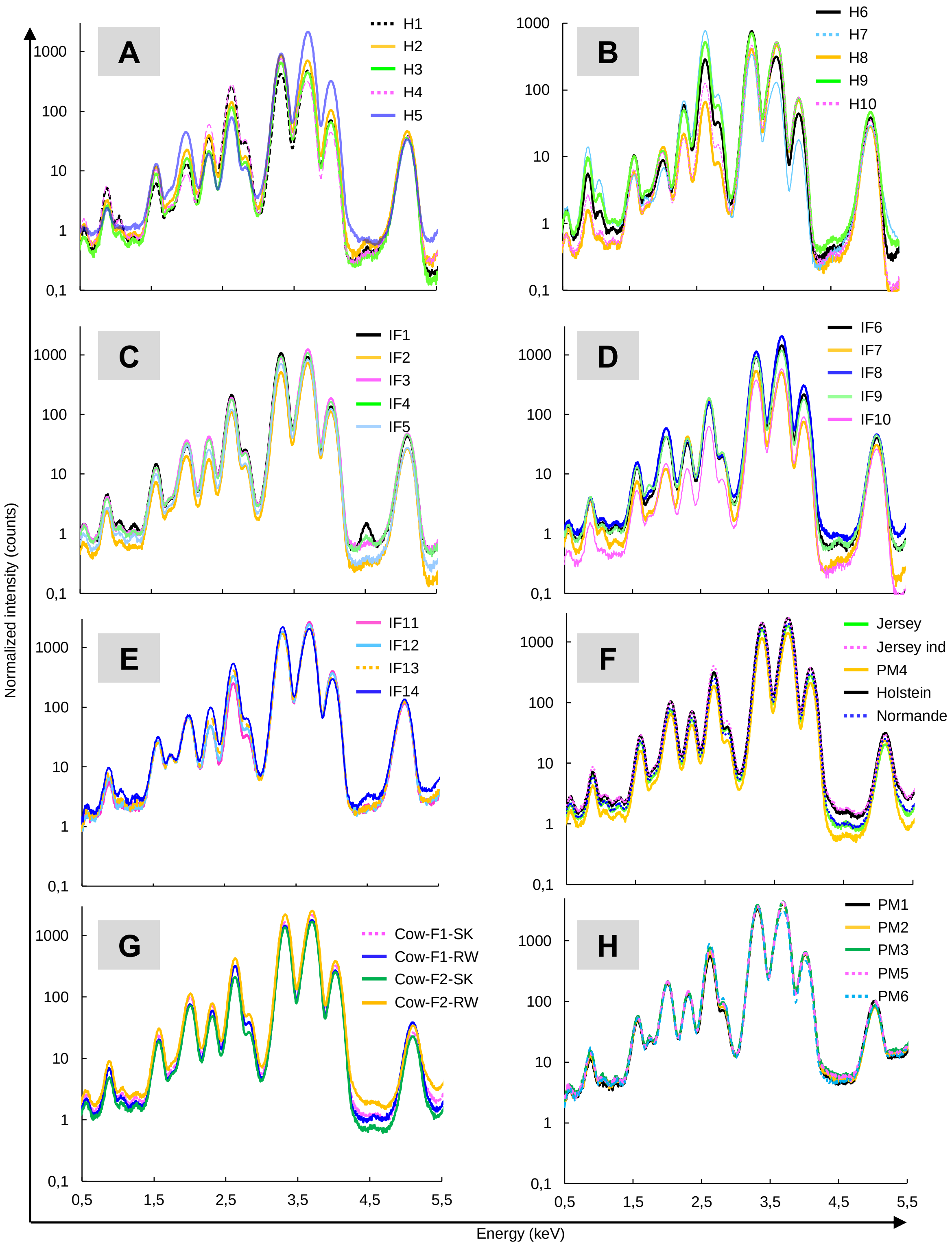

**Fig. S4.** X-ray fluorescence spectra collected at 5.1 keV for IF by brand (A). Spectra of different regions of the same pellet IF3 (B), IF4 (C), IF5 (D), and IF7 (E).

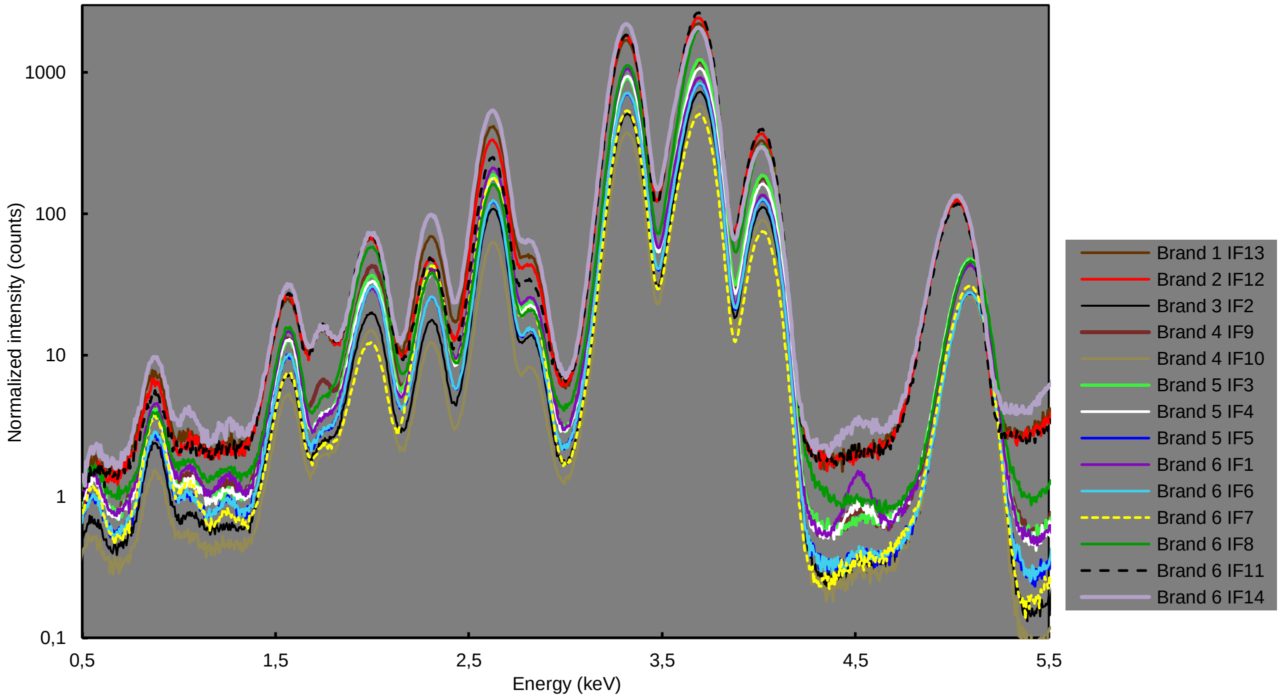

**Fig. S5. Milk production and quality plotted against Ti AUC in the milk of three breeds of cow.** Analysis of milk from Holstein (n=6), Normande (n=6), and Jersey (n=5) cows raised at the INRAE experimental unit. For each breed, individual milk collected on morning milking was blended prior analysis for Ti AUC (red open bars). Ti AUC values were plotted against that of the average per cow of : the volume of milk produced on that day (A), the number of cells per milliliter of milk (B), the protein (C) and fat (D) levels in g per kg of milk.

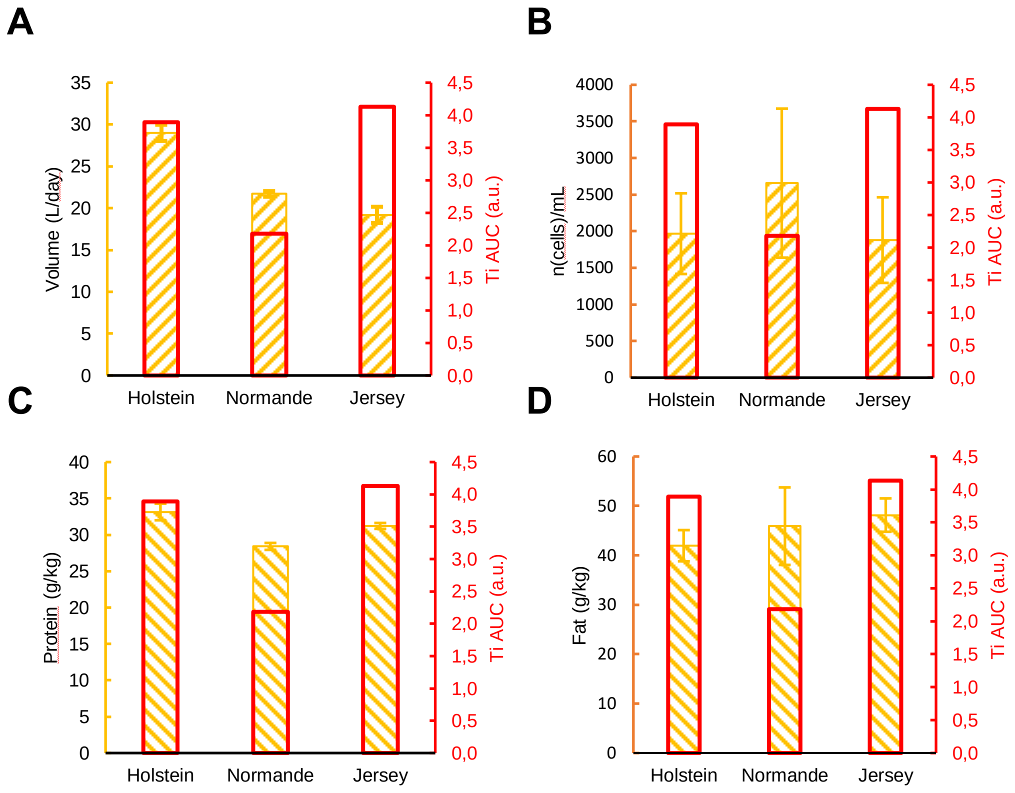

**Fig. S6. Comparison of several regions of the same milk pellet.** (A). X-ray fluorescence spectra collected at 5.1 keV for different regions of the same pellet IF3, IF5, IF7, and IF4. (B) Elements AUC for two regions of IF4.

**A**

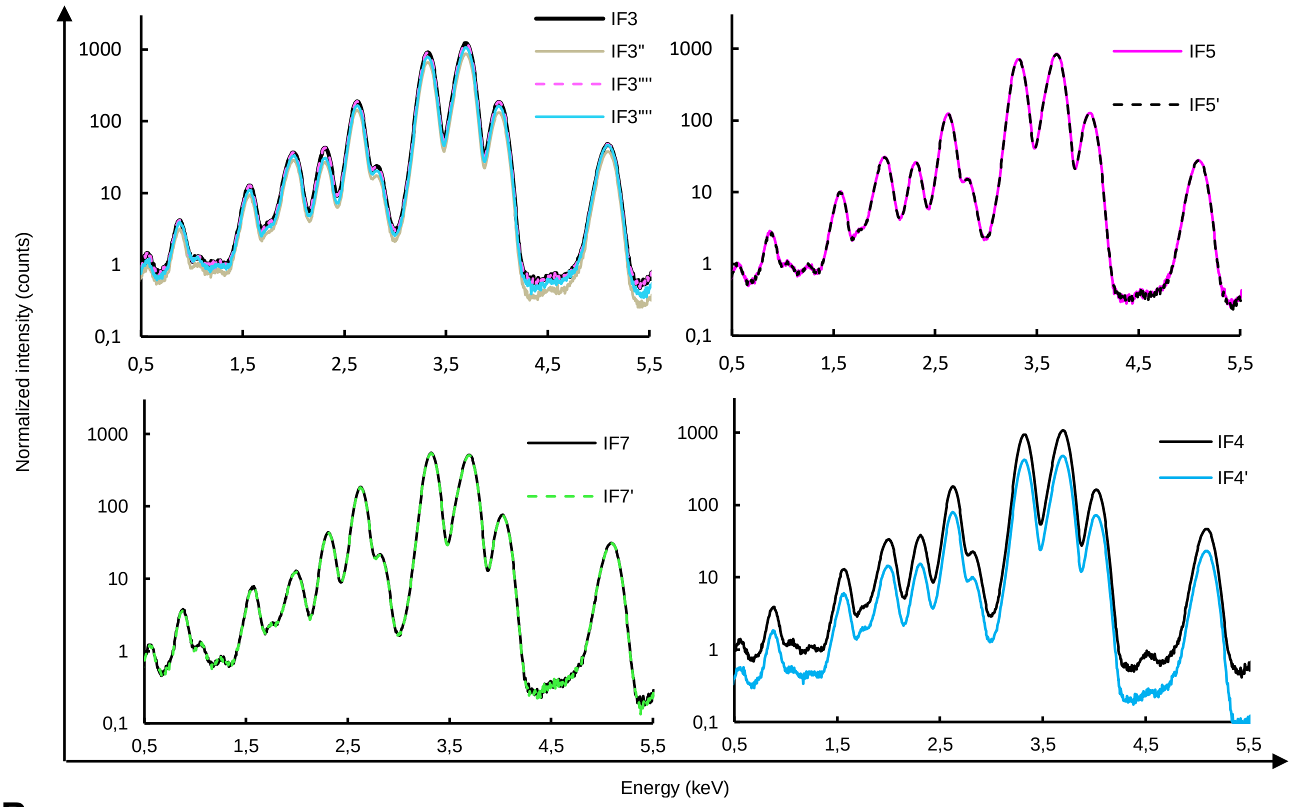


**B**

| | IF4_33470 | IF4_33465 |
|---|---|---|
| O | 2,12 | 4,85 |
| Na | 0,99 | 4,53 |
| Mg | 0,92 | 2,58 |
| Al | 13,96 | 23,42 |
| Si | 8,69 | 8,16 |
| P | 256 | 613 |
| S | 367 | 910 |
| Cl | 2322 | 5288 |
| K | 13519 | 30461 |
| Ca | 16022 | 36209 |
| Ti | 1,09 | 9,48 |
| Cu | 3,49 | 1,92 |

**Fig. S7.** Area under curve (AUC) of Ti and Si peaks computed by PyMCA software.

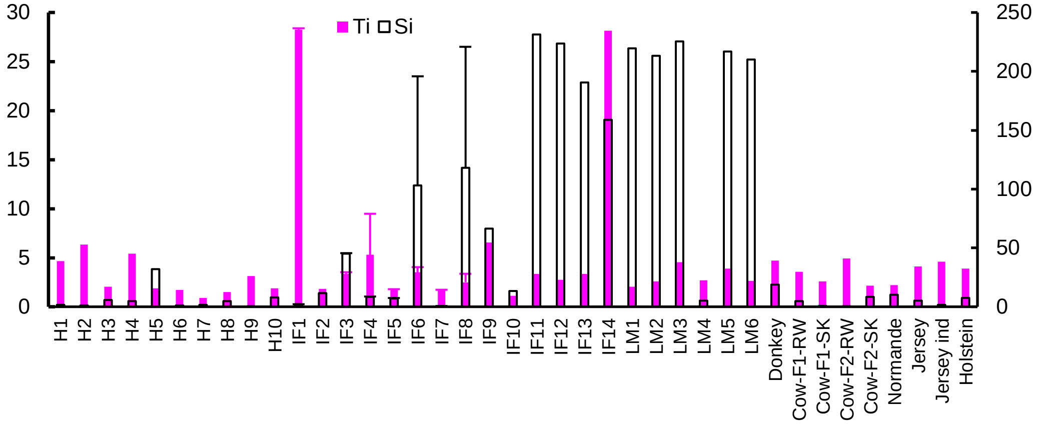


.

**Fig. S8.** Workflow for Ti hotspots quantification on µXRF maps (A). Ti map of human H4 region b (H4b) and the average pixel extracted XRF spectra of 3 regions of interest (ROI): a hotspot (ROI1) and regions without hotspot (ROI 2,3). Black arrows point to Ti peaks (B) Ti maps of a region of H2a before ("raw", left) and after Gaussian filtering (right) on which hotspots were detected automatically (in green). White open arrowheads point to hotspots (C) Number of hotspots (particles) detected for each region

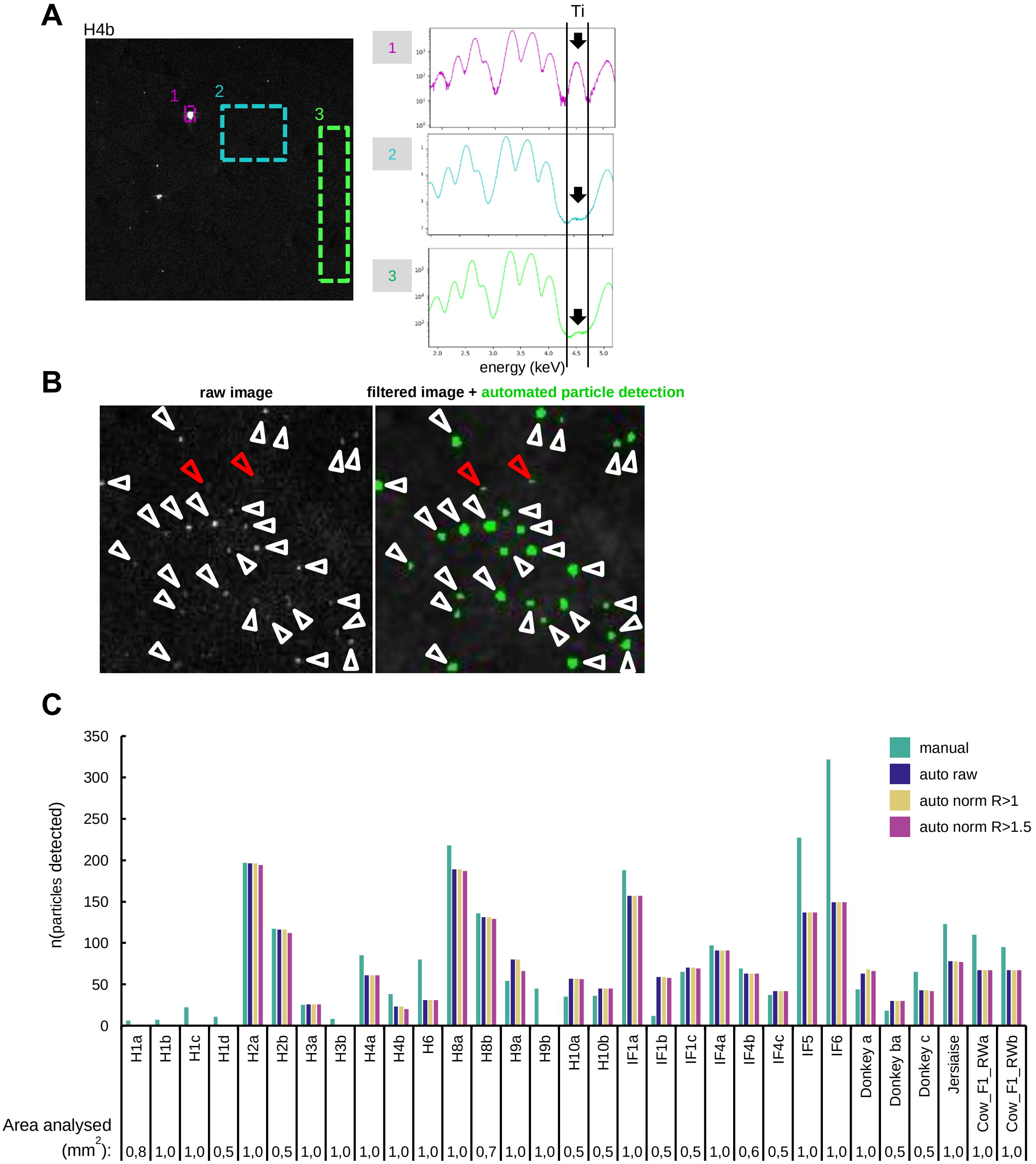


either manually or automatically ("auto", green) without threshold ("auto raw" in blue) or with threshold of 1 (“R>1”, in yellow) or 1.5 (“R>1.5” in pink) signal/noise ratio. Surface area of regions acquired is as indicated.

**Fig. S9.** Ti maps raw ("raw") or Gaussian filtered ("filtered") with automated detected hotspots (in green). Raw histograms of Ti values (left) with the maximum value chosen for display (green graduation).

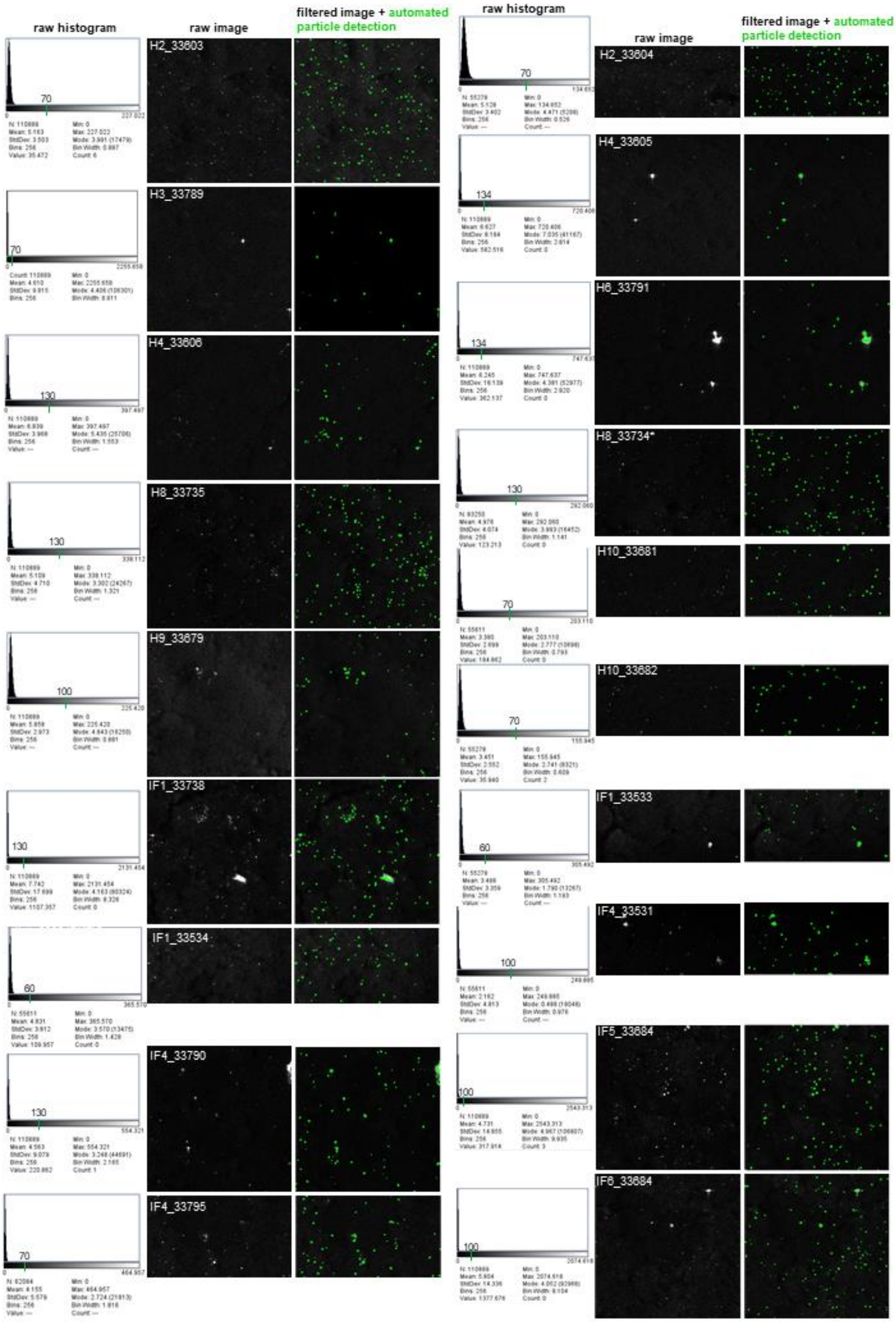

**Fig. S10.** Ti maps raw (“raw”) or Gaussian filtered (“filtered”) with automated detected hotspots (in green). Raw histograms of Ti values are shown (left) with the maximum value chosen for display.

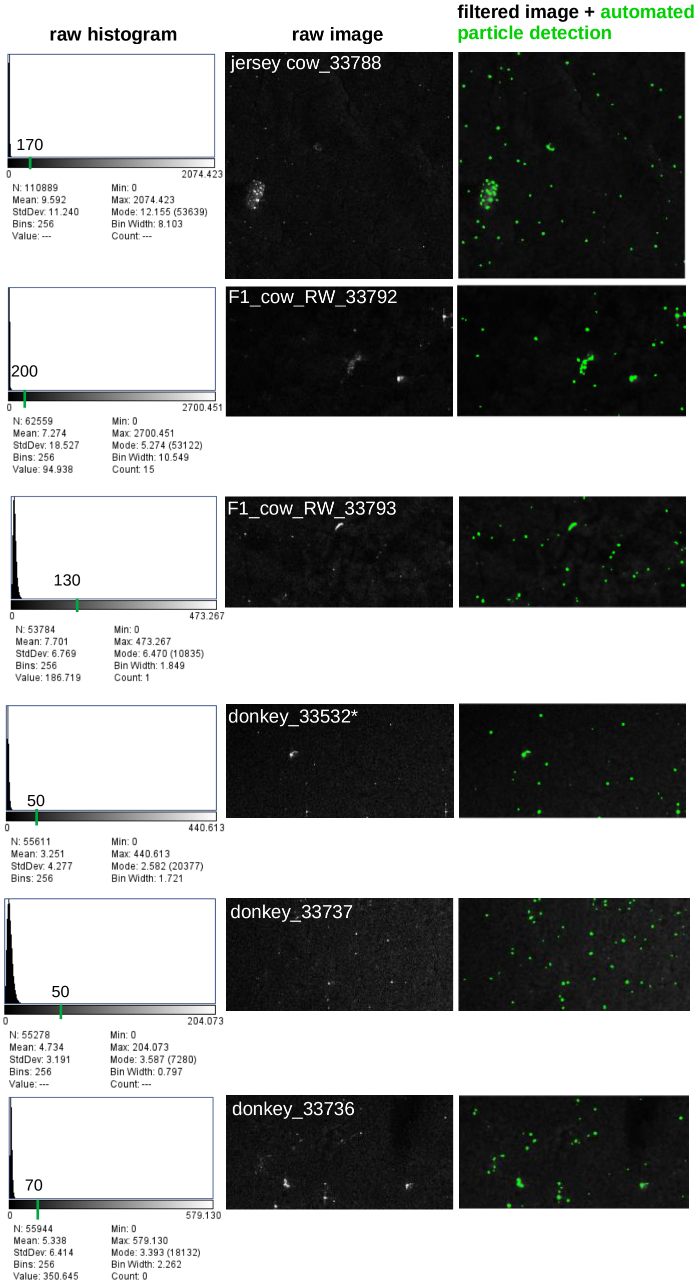

**Fig. S11.** Ti maps raw or Gaussian-filtered with manually detected hotspots (in green). Raw histograms of Ti values is shown (left). Maximal values chosen for display is the maximal value on histogram.

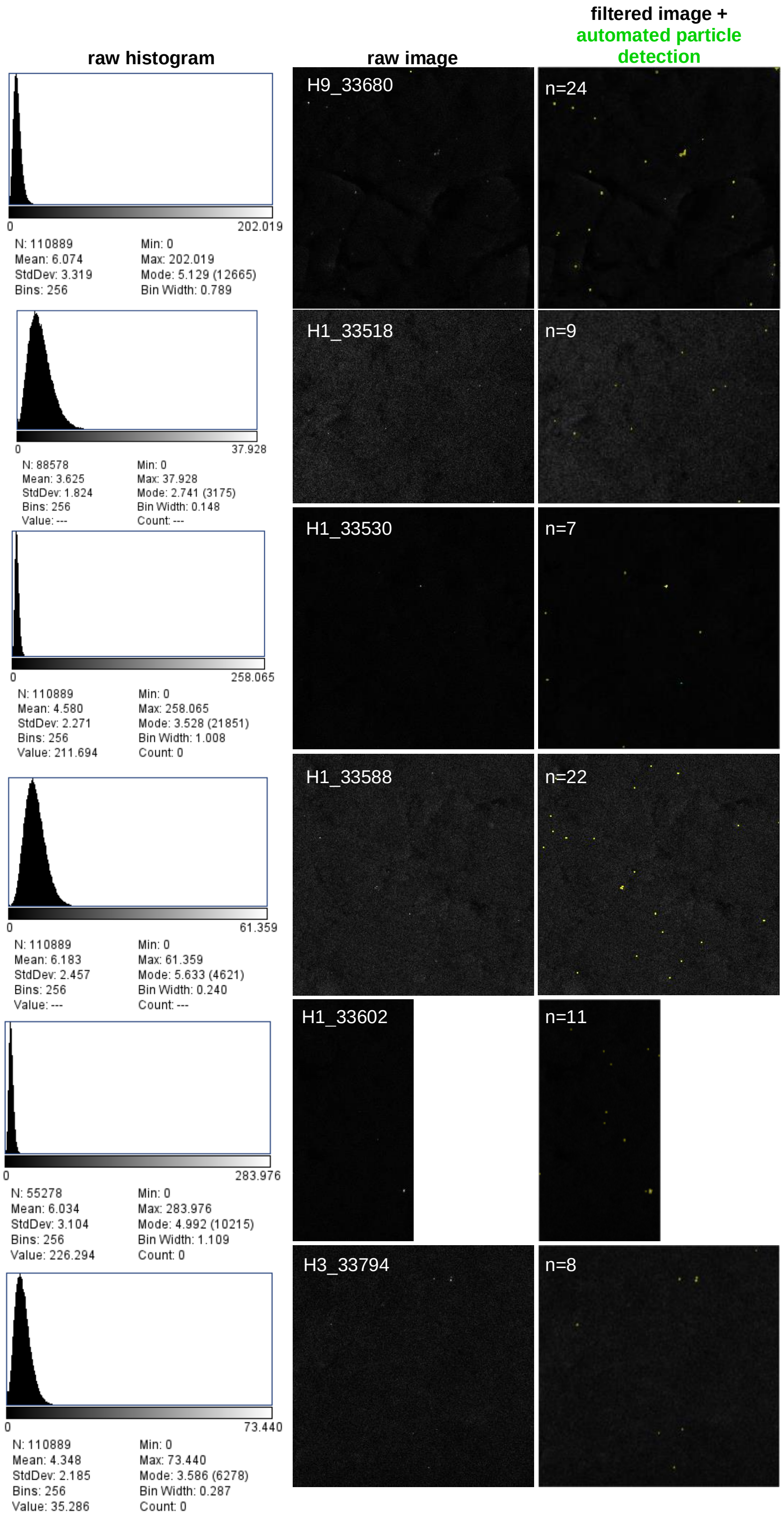

**Figure S12.** Ti K-edge micro-XANES spectra of human (H1, H2, H4, H9, and H10) breastmilk samples and the corresponding attributions of reference spectra (rutile, anatase, ilmenite and pseudobrookite).

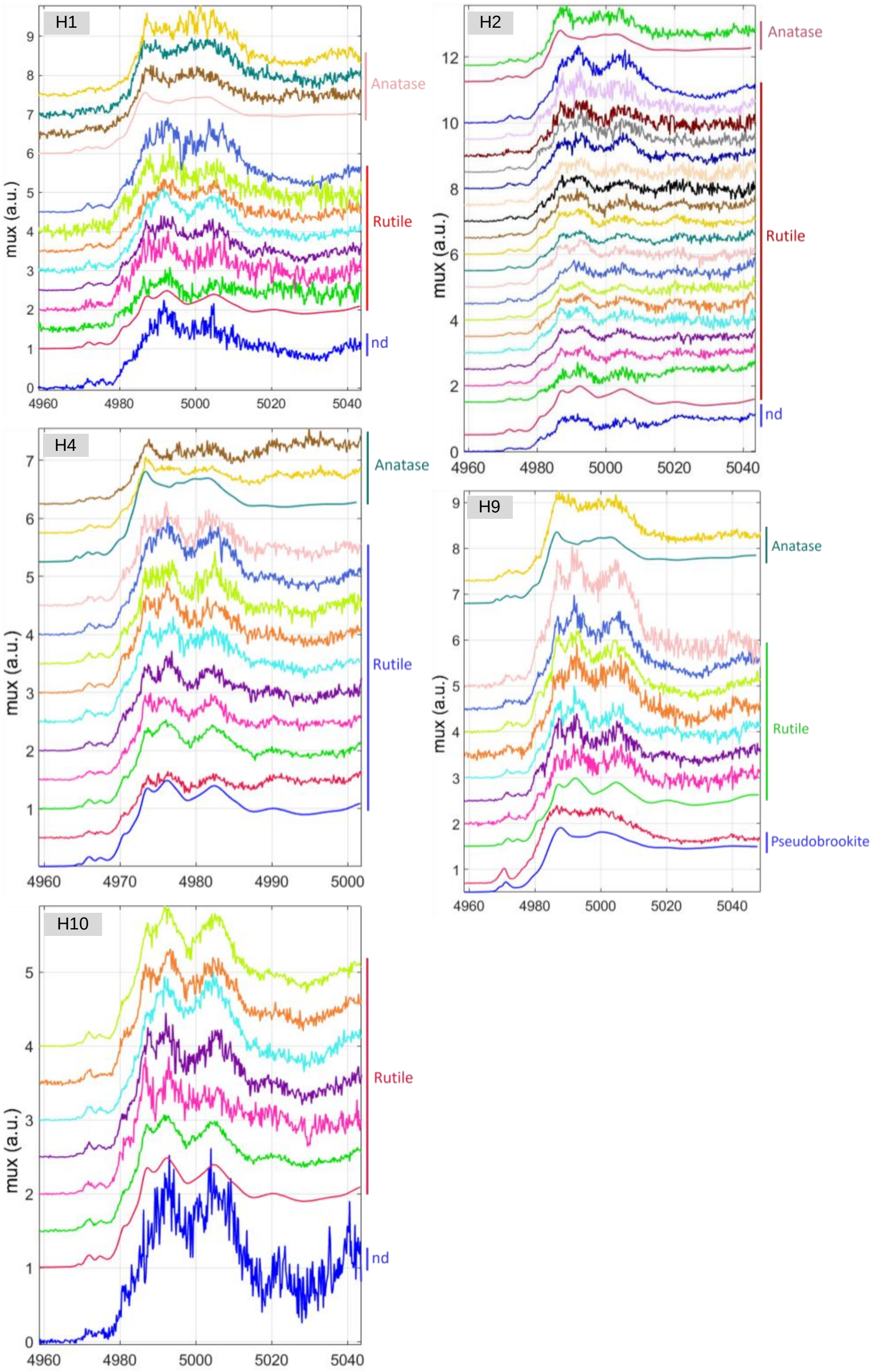

**Fig. S13** Ti K-edge micro-XANES spectra of infant formula (IF1, IF4, IF5, and IF6) and animal milk (donkey and Normande (NOR) cow) and the corresponding attributions of reference spectra (rutile, anatase, ilmenite and pseudobrookite).

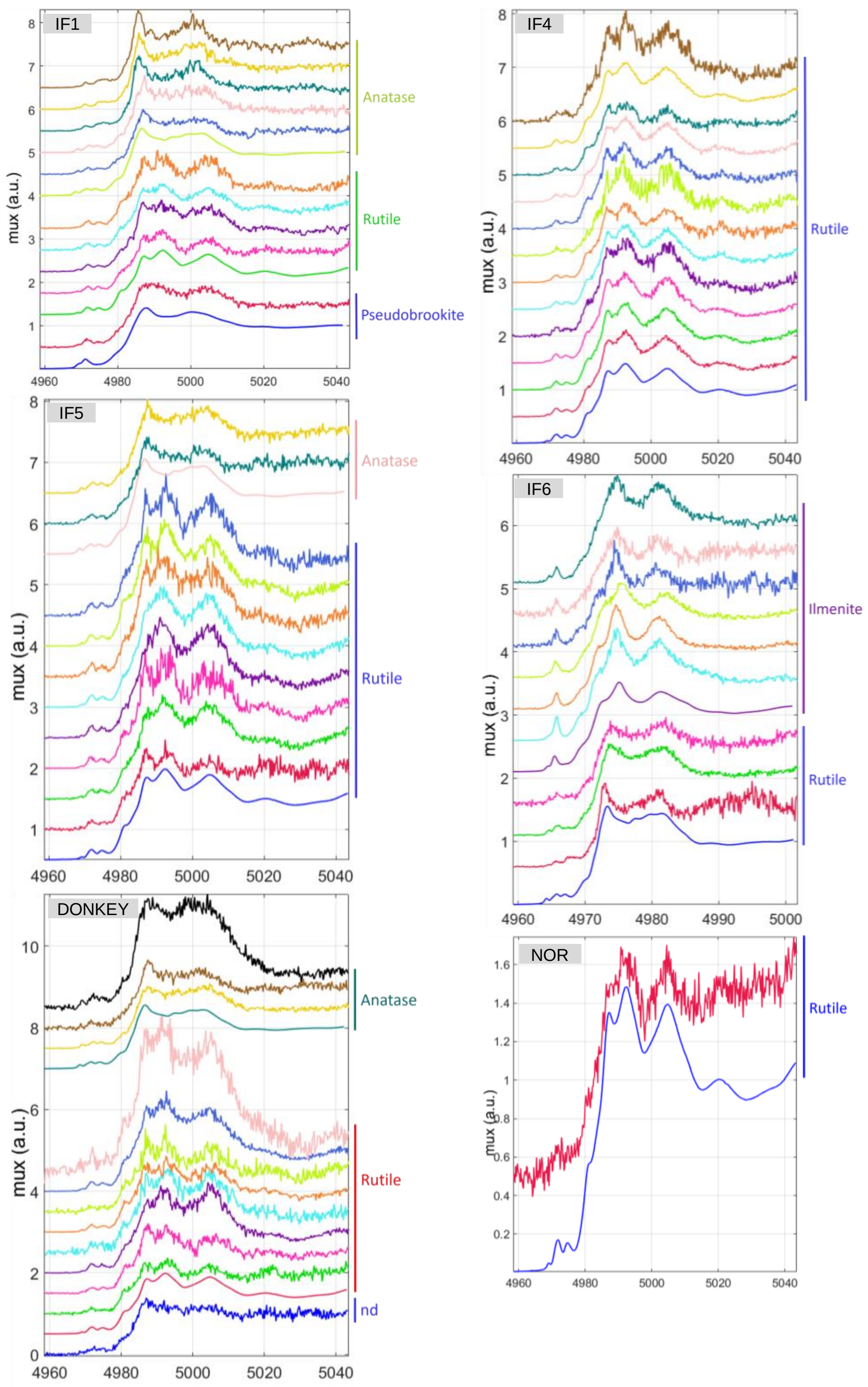

**Fig. S14. Single Particle ICP-MS (SP-ICP-MS) analysis of Ti-containing particles in IF5' reconstituted milk sample**. (A) The entire set of 1 000 000 measures shows high Ti signal peaks corresponding to Ti particles ("particles") and a low continued signal corresponding to soluble Ti ("soluble"). (B) Magnification of the blue dotted box around a single peak.

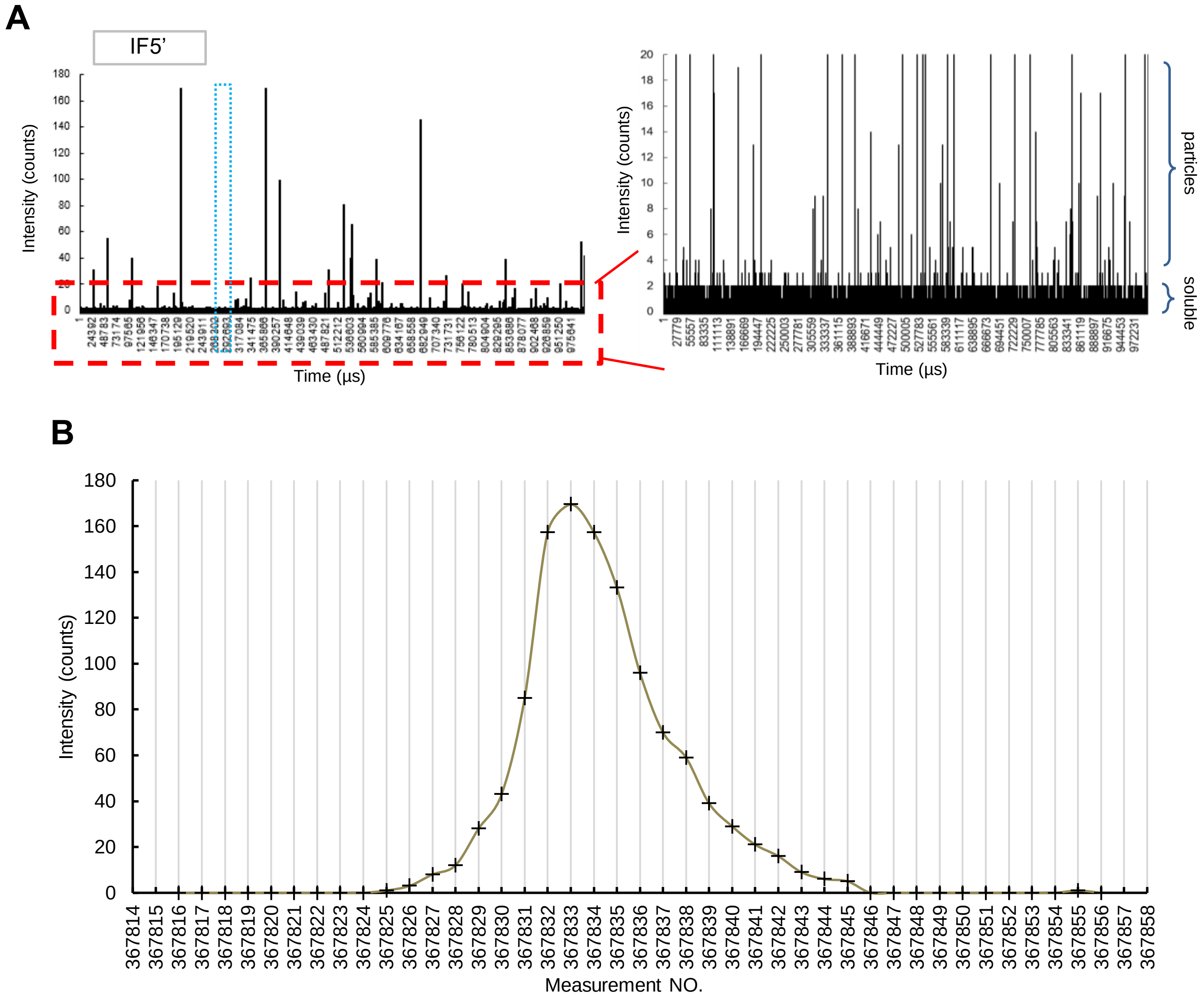

**Fig. S15. Size distribution of Ti-containing particles in milk and NIST standard by SP-ICP-MS.** Frequency (number of particles) and diameter of particles (in nm) are given.

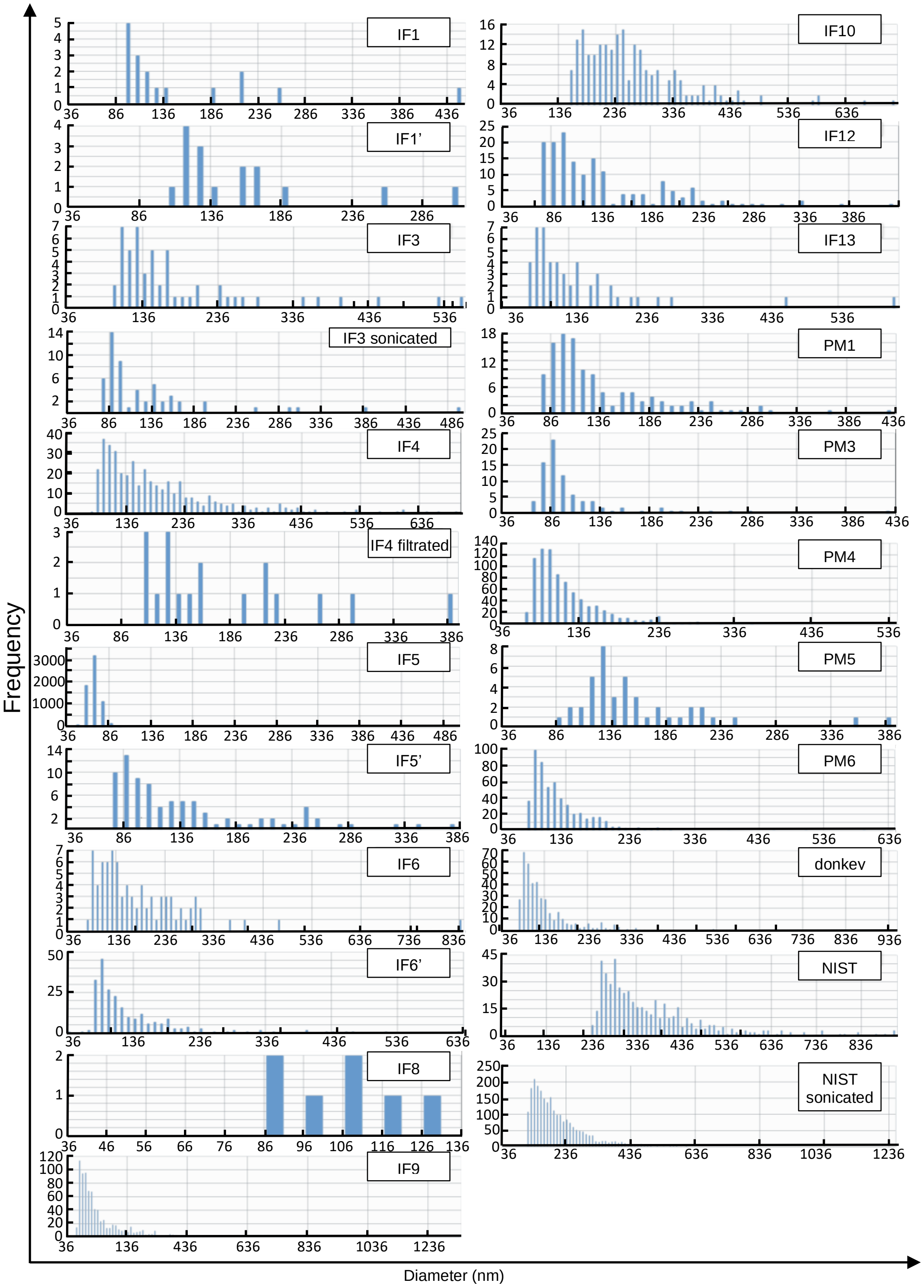

**Fig. S16.** SP-ICP-MS analysis before and after ultrafiltration of IF5’ (A) and ultrapure water (B)

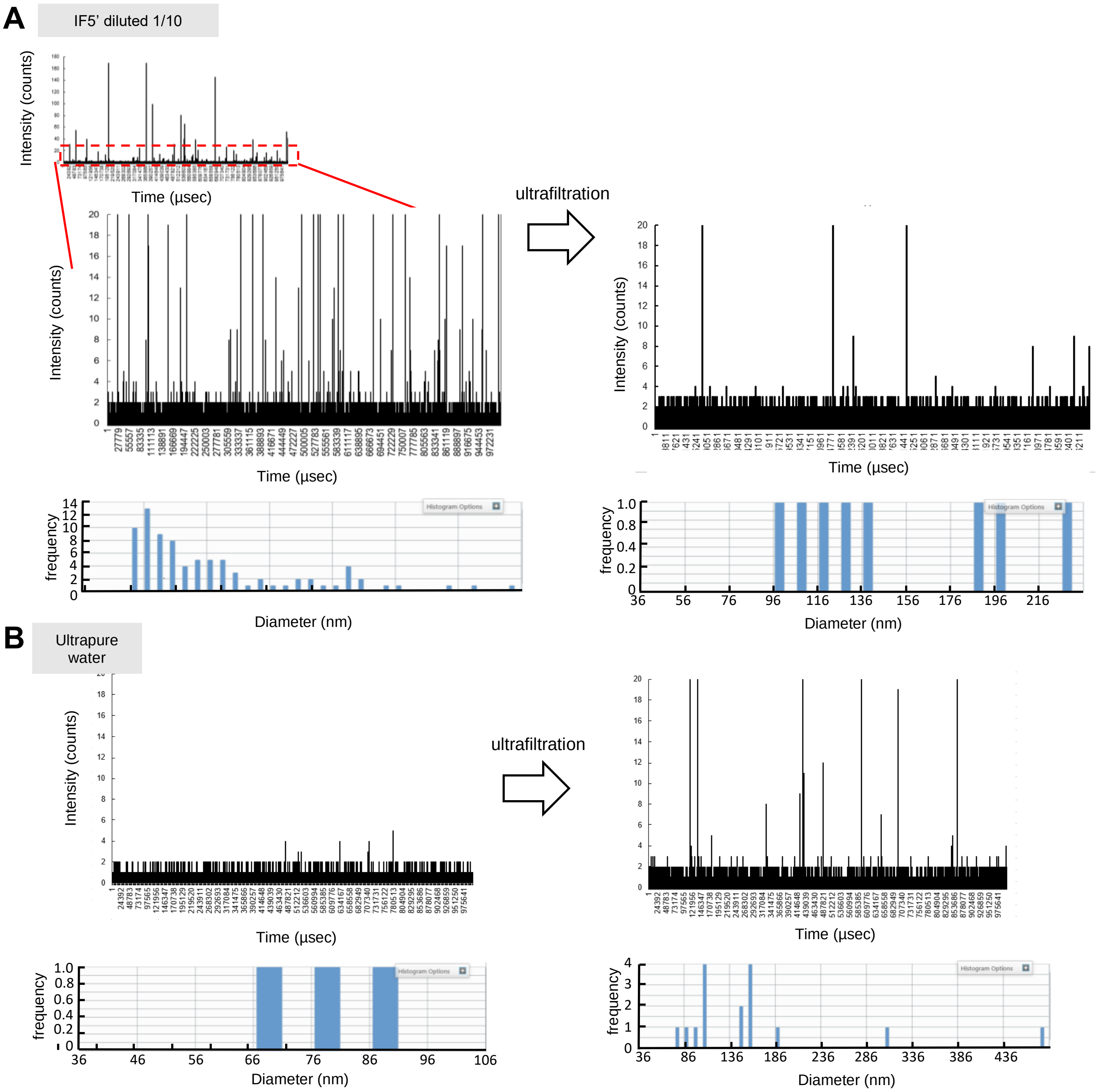

**Table S1** Milk samples type, provider, processing and storage

| ***ID*** | ***type*** | ***provider and origin*** | | ***initial container*** | ***sample processing prior storage*** | ***storage condition*** | ***storage container*** |
|---|---|---|---|---|---|---|---|
| H1 - H10 | human raw | Lactarium Port-Royal (AP-HP) | Paris and Paris region (FR) | bottles CE0459 certified | none | frozen | bottles CE0459 certified |
| IF1-IF14 | infant formula | supermarkets and pharmacies | nd | industrial packaging | none | powder | industrial packaging |
| PM 1-4 | cow powder milk | commercial supermarket | France | industrial packaging | none | powder | falcon 50 Corning polypropylene high clarity/polyethylene cap ISO 13485/9001 |
| PM 5 | cow powder milk | commercial supermarket | Poland | industrial packaging | none | powder | industrial packaging |
| PM 6 | goat powder milk | commercial supermarket | Austria | industrial packaging | none | powder | industrial packaging |
| Donkey | donkey raw milk | farm milk | Ariège county (FR) | glass can | none | frozen | falcon 15 Corning polypropylene high clarity/polyethylene cap ISO 13485/9001 |
| Cow-F1-RW | Mix of breeds, raw tank milk | Farm 1 | Paris region (FR) | Plastic bottle | none | frozen | " |
| Cow-F1-SK | Mix of breeds, skimmed tank milk | Farm 1 | Paris region (FR) | Plastic bottle | skimmed | frozen | " |
| Cow-F2-RW | Mix of breeds, raw tank milk | Farm 2 | Paris region (FR) | Plastic bottle | none | frozen | " |
| Cow-F2-SK | Mix of breeds, skimmed tank milk | Farm 2 | Paris region (FR) | Plastic bottle | skimmed | frozen | " |
| Jersey | Blended milk of 6 Jersey cows collected on day 18/03/2021 | INRAE UEP | Normandie region (FR) | pyrex glass bottle | none | frozen | falcon 50 Corning polypropylene high clarity/polyethylene cap ISO 13485/9001 |
| Jersey ind | raw milk from individual cow collected on day 18/03/2021 | INRAE UEP | Normandie region (FR) | pyrex glass bottle | none | frozen | " |
| Holstein | Blended milk from 6 Holstein cow collected on day 18/03/2021 | INRAE UEP | Normandie region (FR) | pyrex glass bottle | none | frozen | " |
| Normande | Blended miilk of 6 Normande cows collected on day 18/03/2021 | INRAE UEP | Normandie region (FR) | pyrex glass bottle | none | frozen | " |

**Table S2** Additional information on PM and IF samples

| *ID* | *type* | *Information provided by manufacturer* | *Additional information* |
|---|---|---|---|
| PM1 | cow powder milk | half skimmed | |
| PM2 | cow powder milk | half skimmed | |
| PM3 | cow powder milk, organic | half skimmed organic | |
| PM4 | cow powder milk | raw | |
| PM5 | cow powder milk | raw | |
| PM6 | goat powder milk | raw | |
| IF1 | infant formula | 0-6 months old thickened | |
| IF2 | infant formula | 6-12 months old organic | |
| IF3 | infant formula | 0-6 months old | batch 09/2022 |
| IF4 | infant formula | 0-6 months old | IF3 newer batch 06/2023 |
| IF5 | infant formula | 6-12 months old thickened | |
| IF6 | infant formula | 6-12 months old | |
| IF7 | infant formula | 12-36 months old | |
| IF8 | infant formula | 12-36 months old | |
| IF9 | infant formula | 0-6 months old thickened | |
| IF10 | infant formula | 0-6 months old organic | |
| IF11 | infant formula | 12-36 months old organic | |
| IF12 | infant formula | 0-6 months old organic | |
| IF13 | infant formula | 0-6 months old organic | |
| IF14 | infant formula | 0-6 months old thickened | |
| IF1’ | infant formula | 0-6 months old | IF1 new batch 12/2023 |
| IF5’ | infant formula | 6-12 months old | IF5 new batch 12/2023 |
| IF6’ | infant formula | 6-12 months old | IF6 new batch 12/2023 |

**Table S3.** Size and concentration of Ti-containing particles in milk, % of soluble Ti and % of particles as detected by SP-ICP-MS. Concentration is given as Ti particles per liter of milk based on the manufacturer's recommendations to reconstitute powder milk for infant (IF) or adult (PM) consumption.

| *ID* | *Most frequent size(nm)* | *Mean size (nm)* | *Median size (nm)* | *% <100nm* | *Ti part/L* | *% soluble* | *% particles* |
|---|---|---|---|---|---|---|---|
| Water | 86 | 76 | 76 | 100 | 860 000 | | |
| Water filtrated | 116 | 160 | 146 | 19 | 4 610 000 | | |
| IF1 | 96 | 154 | 116 | 29 | 10 290 000 | 97 | 3 |
| IF1' | 116 | 155 | 131 | 0 | 9 681 000 | 96 | 4 |
| IF3 | 126 | 185 | 146 | 4 | 32 067 000 | 61 | 39 |
| IF3 sonicated | 86 | 132 | 96 | 53 | 33 285 000 | 75 | 25 |
| IF4 | 96 | 191 | 161 | 15 | 246 855 000 | 19 | 81 |
| IF4 filtrated | 156 | 178 | 151 | 0 | 10 899 000 | 93 | 7 |
| IF5 | 46 | 66 | 66 | 99 | 3 996 909 000 | 87 | 13 |
| IF5' | 96 | 141 | 116 | 38 | 50 820 000 | 76 | 24 |
| IF5' filtrated | 116 | 149 | 131 | 13 | 4 830 000 | 99 | 1 |
| IF5' sonicated | 66 | 134 | 106 | 42 | 125 853 000 | 52 | 48 |
| IF6 | 126 | 184 | 156 | 14 | 50 211 000 | 86 | 14 |
| IF6' | 56 | 133 | 106 | 46 | 143 997 000 | 84 | 16 |
| IF8 | 96 | 103 | 106 | 43 | 4 830 000 | 99 | 1 |
| IF9 | 216 | 265 | 246 | 0 | 113 631 000 | 26 | 74 |
| IF10 | 46 | 148 | 116 | 40 | 429 030 000 | 28 | 72 |
| IF12 | 96 | 138 | 116 | 39 | 98 616 000 | 79 | 21 |
| IF13 | 86 | 148 | 116 | 31 | 29 652 000 | 85 | 15 |
| PM1 | 96 | 138 | 116 | 34 | 50 414 000 | 79 | 21 |
| PM3 | 56 | 110 | 86 | 66 | 33 474 000 | 89 | 11 |
| PM4 | 46 | 122 | 106 | 46 | 348 096 000 | 76 | 24 |
| PM5 | 126 | 158 | 141 | 7 | 16 940 000 | 95 | 5 |
| PM6 | 56 | 126 | 106 | 41 | 220 640 000 | 42 | 58 |
| Donkey | 96 | 143 | 116 | 36 | 79 971 920 | 42 | 58 |
| NIST 10µg/L | 296 | 365 | 326 | 0 | 146 650 000 | 3 | 97 |
| NIST sonicated 10µg/L | 126 | 214 | 186 | 0 | 705 310 000 | 10 | 90 |
